# DiffModeler: Large Macromolecular Structure Modeling in Low-Resolution Cryo-EM Maps Using Diffusion Model

**DOI:** 10.1101/2024.01.20.576370

**Authors:** Xiao Wang, Han Zhu, Genki Terashi, Manav Taluja, Daisuke Kihara

## Abstract

Cryogenic electron microscopy (cryo-EM) has now been widely used for determining multi-chain protein complexes. However, modeling a complex structure is challenging particularly when the map resolution is low, typically in the intermediate resolution range of 5 to 10 Å. Within this resolution range, even accurate structure fitting is difficult, let alone de novo modeling. To address this challenge, here we present DiffModeler, a fully automated method for modeling protein complex structures. DiffModeler employs a diffusion model for backbone tracing and integrates AlphaFold2-predicted single-chain structures for structure fitting. Extensive testing on cryo-EM maps at intermediate resolutions demonstrates the exceptional accuracy of DiffModeler in structure modeling, achieving an average TM-Score of 0.92, surpassing existing methodologies significantly. Notably, DiffModeler successfully modeled a protein complex composed of 47 chains and 13,462 residues, achieving a high TM-Score of 0.94. Further benchmarking at low resolutions (10-20 Å) confirms its versatility, demonstrating plausible performance. Moreover, when coupled with CryoREAD, DiffModeler excels in constructing protein-DNA/RNA complex structures for near-atomic resolution maps (0-5 Å), showcasing state-of-the-art performance with average TM-Scores of 0.88 and 0.91 across two datasets.

## Introduction

Proteins are fundamental molecules that carry out numerous functions in living organisms, including enzyme catalysis, cell signaling, and transport of molecules Cryogenic electron microscopy (cryo-EM) has gained significant popularity among experimental protein structure determination techniques^1–3^. This technique is increasingly favored due to several advantages, notably its superior capacity to determine the three-dimensional (3D) structures of large macromolecular complexes.

While reported map resolutions in literature have generally shown steady improvement over recent years, it remains common to encounter intermediate resolutions (∼5-10 Å) in real-life lab scenarios, posing challenges for structure modeling. When the map resolution is better than 5 Å, direct tracing of main-chain of proteins^4–8^ and nucleic acids^9^ have now become feasible due to recent modeling methods leveraging deep learning to detect atom positions within the map. However, for maps within the intermediate resolution range (5-10 Å), de novo modeling is generally not viable because the identification of amino acid residues and atoms remains elusive even with deep learning techniques. Hence, a practical approach involves conducting structure fitting using methods such as Phenix^10^, Flex-EM^11^, Assembline^12^, MultiFit^13^, Chimera^14^, MarkovFit^15^, and VESPER^16^ or employing manual fitting with known structures from PDB^17^ or predicted structure models^18^. Secondary structure detection methods within EM maps^19^ ^20,21^ can aid in protein structure fitting. Despite many structures are determined through structure fitting, accurately orienting molecules within a noisy map in this resolution range remains challenging, especially for complexes comprising multiple subunits. The successful development of an automatic and precise structure fitting method for EM maps at medium- and low-resolution intermediate resolutions would significantly support structural biologists.

Here, we developed DiffModeler, a fully automated structure fitting method for modeling large protein complex structures in cryo-EM maps with resolutions ranging from 5 to 10 Å. DiffModer uses a diffusion model^22–24^ to enhance the map aiding in finding precise fitting poses for these structures. The diffusion model is a parameterized Markov chain trained using variational inference to generate samples that match the underlined data after a finite time frame. Notably, the diffusion model has demonstrated considerable success in various areas of image processing, such as image generation^23–26^, segmentation^27,28^, and translation^29,30^. Additionally, it has found applications in bioinformatics, including protein docking^31^ and protein design^32,33^. Building upon these successful applications, DiffModeler integrates the diffusion model to enhance the extraction of structural information, facilitating accurate structure modeling for cryo-EM maps at intermediate resolutions.

To the best of our knowledge, this is the first fully automated and accurate method for modeling protein complex structures in maps at intermediate resolutions. DiffModeler initiates the process by tracing protein backbones within a cryo-EM map, employing a diffusion model designed to capture the distinctive local density patterns representing protein backbones. Simultaneously, we use AlphaFold2 (AF2) ^34^, the cutting-edge protein structure prediction method, to generate high-quality single-chain structures. Subsequently, the structure models from AF2 are fitted into the traced backbone map, producing many candidate poses through the VESPER^16^ structure fitting program. Ultimately, the complete protein complex structure is assembled by combining candidate poses of constituent subunits.

A benchmark conducted on 19 EM maps ranging from 5.0 Å to 10.0 Å resolution, depicting large protein complexes of 3 to 48 subunits, demonstrated that modeling with DiffModeler substantially outperformed conventional methods^10,16^. On average, the models by DiffModeler had a TM-score^35^ of 0.92, a significant contrast to the conventional methods that yielded an average TM-score of 0.41. Extending our evaluation, we further benchmarked DiffModeler on 6 experimental maps at a low resolution of 10 to 20 Å, where DiffModeler modelled the structure with a TM-Score of 0.27 to 0.97. Additionally, we integrated DiffModeler with CryoREAD, our DNA/RNA structure modeling method^9^, to build protein-nucleic acid complex structures in two datasets comprising 61 and 28 maps at a resolution of up to 5 Å. This combined protocol showcased a state-of-the-art performance delivering an average TM-Score of 0.88 and 0.91, respectively.

## Results

### Overview of DiffModeler

We begin by explaining the DiffModeler algorithm depicted in Fig .1. DiffModeler comprises four major steps: First, it detects the protein backbone positions in the input cryo-EM map by enhancing the map using a trained diffusion model. Secondly, it conducts the modeling of individual protein structures using AF2. Thirdly, structure models are fitted to the enhanced map using VESPER. Lastly, it selects and combines fitted single-chain poses to build the complete protein complex structures within the map. Below, we provide more information of each step.

### Backbone Tracing via Diffusion Model

Achieving accurate structure fitting for maps of an intermediate resolution is untrivial. To aim for higher accuracy, the main innovation of DiffModeler is to use diffusion model to pronounce the density that belong to protein backbone. The input map is scanned with a 64^3^ Å^3^ box with a stride of 32 Å along the map grid with a 1 Å interval. Given a box of cryo-EM density, the encoder of the conditional diffusion model computes an embedding of the input density box. Subsequently, the decoder starts with random Gaussian noise as the initial density distribution and iteratively refines its estimates to make it closer to the ground-truth traced backbone conditioned on the embedding from the encoder and the initial density input. The diffusion process is illustrated in Fig. 1b. This traced backbone provides clearer information for structure fitting compared to the original map. The diffusion model is trained via denoising diffusion implicit model (DDIM) framework. During training, the main objective of the model is to perform conditional denoising of a noisy density of the traced backbone to achieve the ground-truth traced protein backbone density in the map. The overall framework is optimized via Dice loss^36^ that considers the agreement of the identified and ground-truth backbone positions. The training and inference framework is presented in Extended Data 1 and Extended Data 4, respectively, and further details regarding training and inferences can be found in Methods.

**Fig. 1.**
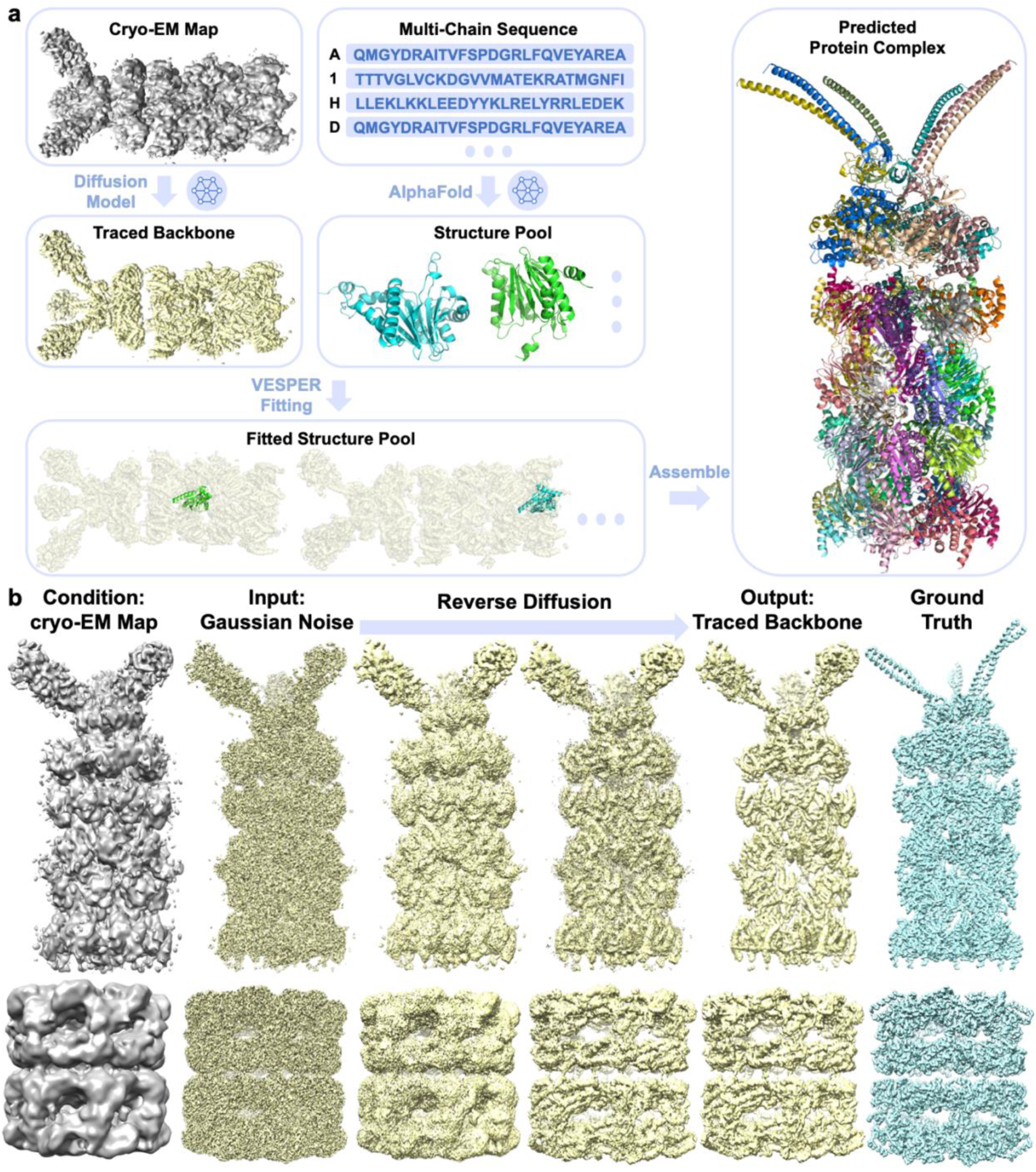
Overall framework of DiffModeler. **a.** Workflow of DiffModeler. DiffModeler consists of four main steps. 1) Backbone tracing from cryo-EM maps at intermediate resolution via a diffusion model. 2) Single-chain structure prediction by AF2. 3) Single-chain structure fitting using VESPER. 4) Protein complex modeling by assembling algorithms. **b.** Overview of Diffusion Process. Starting from an original cryo-EM map as a condition and random Gaussian noise. The protein backbone is traced by the iterative reverse diffusion process utilizing a pre-trained diffusion model. On the right shown is the ground-truth protein backbone density, which is the target of the reverse diffusion process. The examples used here are EMD-0213 (Resolution: 6.35 Å) and EMD-1042 (Resolution: 10.3 Å).

### Structure Prediction by AF2

In DiffModeler, we use predicted single chain protein structures by AF2^34^ to fit into the diffused map of pronounced backbone structure. While there are instances where AF2 models do not align with the proteins’ conformations in particular cryo-EM maps^37^, ample cases exist^38–40^ where AF2 models demonstrated sufficient accuracy to be effectively integrated into EM maps. Specifically, for maps at a resolution worse than 5 Å, the resolution range of the main DiffModeler’s objective, where de novo main-chain tracing becomes highly challenging, it would be pragmatic to consider AF2 models for structure modeling. Instead of generating new AF2 models, users can also use precomputed models available in the AlphaFold database^18^, which we employed in this work.

### Structure Model Fitting with VESPER

The predicted structure models are fit to the diffused map using VESPER^16^, a structure and map fitting method developed in our group. Unlike the conventional methods that solely rely on correlation of map densities, VESPER takes into account local density gradient within maps. This approach has demonstrated superior performance surpassing existing methods^16^. The predicted structure models are converted into simulated maps at a 1 Å resolution. Subsequently, both these simulated maps of derived from the models and the diffused main-chain map are transformed into local dense points (LDPs) using the mean-shift algorithm^4,41^. LDPs serve to encapsulate the local salient features of density, proving to be more precise for alignment than using the unprocessed maps alone. Using VESPER, each subunit is aligned with the diffused main-chain map, resulting in the retention of the top 100 candidate poses.

### Protein complex modeling by a greedy assembling algorithm

This phase is geared towards assembling the complete protein complex structure through an assembly of suitable poses from each subunit. To accomplish this, we have devised a greedy algorithm, which is explained in detail in the Methods section and visually outlined in Extended Data 5. In the preceding step, a collection of 100 poses has been constructed for each subunit, with each pose being evaluated based on a fitness score. From all combinations of subunit-pose pairs, we identify the subunit-pose with the highest score. Subsequently, we mask the local density of the map occupied by this selected subunit-pose pair and select the next best subunit-pose in the pool. This process iterates, systematically selecting the subsequent best subunit-pose pairs until all subunits seamlessly integrate into the emphasized diffused protein backbone map.

### Structure Modeling Performance

We assessed DiffModeler’s performance on an independent dataset comprising 19 maps determined at resolutions between 5.0 Å and 10.0 Å. These structures are nonredundant in comparison to the training and validation datasets we used (see Methods). Supplementary Table 1 provides a comprehensive list of the maps included in this dataset. The range of residues of proteins in these maps varied from 1,202 to 13,462, while the number of protein chains ranged between 3 and 47. Notably, 12 out of 19 maps include protein complexes with more than 3,000 residues in total, which is larger than the size that the state-of-the-art protein docking method, Alphafold-Multimer^42^ was trained on and the size that Alphafold Colab notebook can handle.

Fig. 2 summarizes the modeling accuracy on the dataset from various perspectives. In Fig. 2a, we assessed the accuracy of the diffused backbone map generated by the diffusion model in the initial step of DiffModeler (as depicted in the traced backbone panel in Fig. 1). We computed recall and precision of the grid points within the diffused map with reference to the backbone heavy atoms (Cα, C, N) excluding oxygen of proteins in a map. The backbone recall was computed for each residue by determining the fraction of backbone heavy atoms within a 3 Å proximity to any grid points in the diffused map. This was then averaged across all residues in the map. Precision, on the other hand, was computed as the fraction of grid points within a 3 Å proximity to any backbone atoms. As a diffused map outlines backbone atom positions within an input EM map, the volume of a map was, in principle, reduced, on average, by 53.7%. This modification of maps notably elevated the average precision to 85.1% from 68.8% without significantly compromising the recall, which remained stable at an average of 93.1% from 96.6% (the original maps). Detailed results of individual maps are available in Supplementary Table 2.

**Fig 2.**
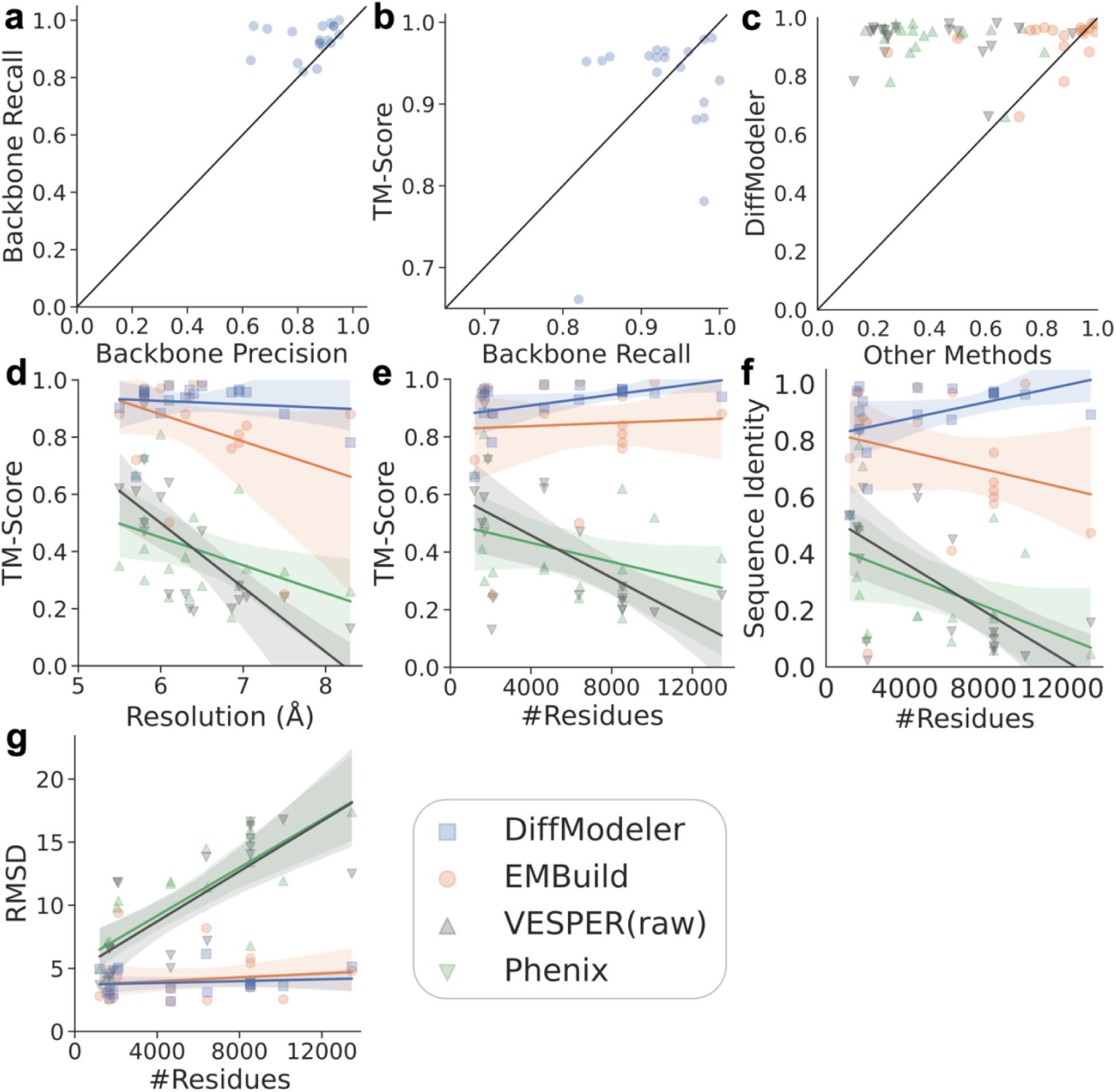
Performance of protein complex structure modeling by DiffModeler. **a.** Backbone recall and precision of the diffusion model. Recall and precision were computed by considering grid points in the diffused maps relative to ground-truth positions of main-chain atoms in the maps. Details in Supplementary Table 2. **b.** TM-Score of modeled protein complex structure relative to backbone recall. **c.** TM-Score comparison between DiffModeler and three existing methods, the raw VESPER, Phenix, and EMBuild. **d.** TM-Score relative to the map resolution of the different methods. **e.** TM-Score relative to the overall protein complex size represented by the number of residues for the different methods. **f.** Sequence identity relative to the structure size for the different methods. **g.** RMSD relative to the structure size for the different methods. The lines in the panels represent the regression line and the shaded region represents the confidence interval for the regression estimate. Raw data of the plots are available in Supplementary Table 3.

Fig. 2b illustrates our exploration into the impact of backbone recall within diffused maps on the subsequent accuracy of structure fitting. To assess the precision of modeled protein complexes, we employed MM-align^43^ to superimpose a modeled complex structure onto the accurate structure (referencing the PDB entry of the complex associated with the EMDB entry of the map) and calculated the TM-Score^35^. The TM-Score is a dimensionless metric utilized to gauge structural resemblance between two protein structures, with a value of 1 denoting identical protein pairs and values exceeding 0.5 indicative of meaningful similarity. Across the entire spectrum of observed backbone recall within this dataset, the TM-Score of modeled structures consistently registered notably high values, exceeding 0.9 for the majority of cases. On average, DiffModeler achieved a high TM-Score of 0.808. There were two instances where the TM-Score fell below 0.8. Notably, in one particular case (EMD-1871), despite a high backbone recall of 0.98 (close to 1.0), the TM-Score remained 0.781 (close to 0.8). This occurred as a result of two wrong single-chain structure fittings because of the low backbone precision 0.64.

In Fig. 2c, we compared the TM-Score of models constructed by DiffModeler with three other existing methods, the *dock_in_map* program in Phenix^10^, EMBuild^44^, and structure fitting by the raw VESPER^16^. For the latter, the original EM maps were used instead of the diffused maps for structure fitting. EMBuild is a recent method for fitting AF2 models within a cryo-EM map, which combines structure fitting, domain-based refinement, and graph-based iterative assembly. For all cases except two (or one) against Phenix, DiffModeler exhibited significantly higher TM-Scores compared to the three methods. The models produced by DiffModeler demonstrated an average TM-Score of 0.922. In contrast, VESPER (raw), Phenix, and EMBuild showcased a broad spectrum of model accuracy, averaging approximately half of DiffModeler’s performance, with TM-Scores of 0.407, 0.409, and 0.841, respectively. The notable contrast between DiffModeler and VESPER (raw) vividly highlights the substantial positive impact of utilizing diffused maps.

Figs. 2d to 2g aim to explore the relationship between model accuracy and both map resolution (Fig. 2d) and the size of the complexes (Fig. 2e, 2f, and 2g). While the performance of other methods noticeably declined with increasing resolution and larger structure sizes, DiffModeler consistently maintained stable performance and notably outperformed in challenging scenarios involving lower resolutions or larger sizes. Fig. 2f compares the sequence identity of different methods relative to the structure size, which considers the fraction of residues in the reference structure that were successfully modeled and with the correct residue type. DiffModeler demonstrated the stable sequence identity, while all other methods decreased dramatically when the structure size was large. On average, DiffModeler, VESPER (raw), Phenix, and EMBuild yielded sequence identities of 0.89, 0.31, 0.29, and 0.74. respectively. Fig. 2g investigates model accuracy concerning complex sizes using a different metric, the root-mean-standard-deviation (RMSD) of the aligned residues in the model, defined as the residues that are aligned by MM-Align’s superimposition. On average, DiffModeler, VESPER (raw), Phenix, and EMBuild yielded RMSD values of 3.89 Å, 10.09 Å, 10.48 Å, and 4.08 Å, respectively. Supplementary Table 3 provides the results of individual maps for different methods.

### Examples of Protein Complex Structure Models

In this section we discuss five examples of models constructed by DiffModeler. In Fig. 3, for each example map, five panels are shown: the original experimental map, the diffused backbone map, LDPs of traced backbone, the structure models, and structure comparison between the constructed model with the PDB entry. The first example (Fig. 3a) is the state 2 of *Mus musculus* TRPML1 (EMD-6824, resolution: 7.4 Å), which encompasses four protein chains totaling 1,696 residues^45^. The resolution of this map was mentioned to be 7.4 Å in the paper^45^ but it may be even worse because Resmap^46^, a map resolution estimation program, reported 9.4 Å when we ran it. Modeling the interaction between the transmembrane domain and the peripheral domain was particularly difficult for this map, resulting in low TM-Scores of 0.30 and 0.47 using Phenix and VESPER (raw), respectively. In contrast, DiffModeler nicely traced the backbone by diffusion model. Thus, it demonstrates remarkable accuracy for structure modeling, achieving a TM-Score of 0.95 and an align ratio of 1.0, thereby providing a very precise structure for this challenging map.

**Fig. 3.**
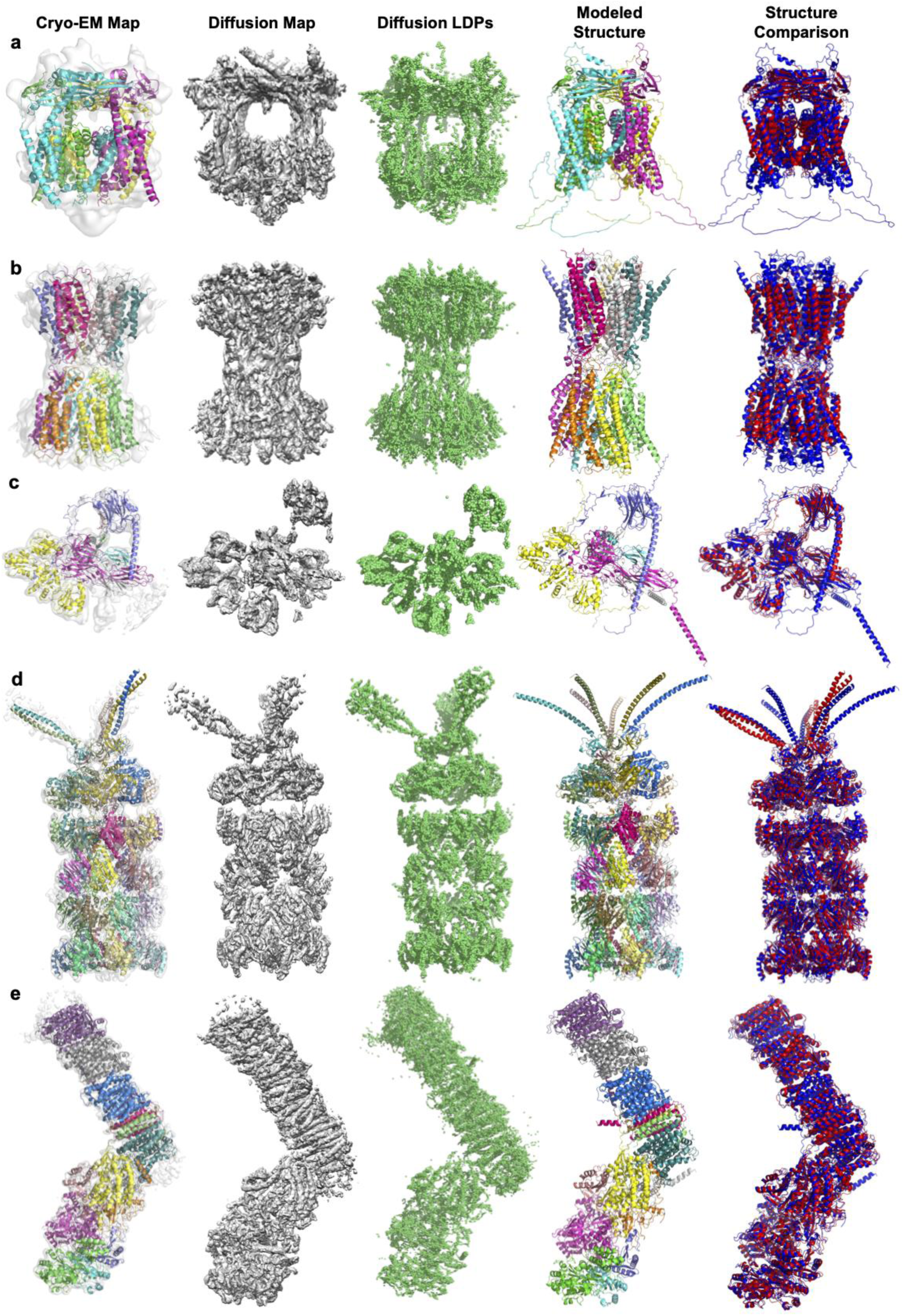
Examples of structure models constructed by DiffModeler from the test dataset. Detailed evaluation Results are available in Supplementary Table 3. For each example, five columns are shown: the cryo-EM map with the structure of the protein complex with different color indicating different chains; the diffused backbone map by the diffusion model; the local dense points of the backbone map; the structure model by DiffModeler; the superposition of the model by DiffModeler (blue) with the native structure (red). **A.** State 2 of *Mus musculus* TRPML1 (EMD-6824, PDB: 5YE1, Resolution: 7.4 Å (see the text); protein size: 4 chains and 1,696 amino acids (aa)). TM-Score: 0.95; RMSD: 3.26 Å. **b.** Closed conformation of Cx26 Gap junction channels at acidic pH (EMD-20916, PDB: 6UVT, Res.: 7.50 Å; protein size: 12 chains and 2,112 aa). TM-Score: 0.88; RMSD: 5.04 Å. **c.** the human PLC editing module (EMD-3906, PDB: 6ENY, Res.: 5.80 Å; protein size: 5 chains and 1,569 aa). TM-Score: 0.95; RMSD: 3.08 Å. **d.** State 2 of ATPase cycle in PAN-proteasomes (EMD-213, PDB: 6HE9, Res.: 6.35 Å; protein size: 34 chains and 8,531 aa). TM-Score: 0.97; RMSD: 3.79 Å. **e.** Minor state of T. *thermophilus* enzyme in complex with NADH (EMD-11237, PDB: 6ZJN, Res.: 6.10 Å; protein size: 15 chains and 4,655 aa). TM-Score: 0.98; RMSD: 2.40 Å.

The next example (Fig. 3b) is the closed conformation of Cx26 Gap junction channels (GJCs) at acidic pH (EMD-20916)^47^. This complex is difficult to model because it has 12 chains in a map of a relatively low resolution, 7.5 Å. DiffModeler was able to precisely identify helices in the map and correctly fit the 12 chains with a TM-Score of 0.88. In contrast, VESPER (raw) struggled to find correct poses of the chains, resulting in a TM-Score of 0.24.

Fig. 3c is the model for the human peptide-loading complex (PLC) editing module (EMD-3906, resolution: 5.8 Å)^48^. Modeling the full protein complex is difficult due to the substantial flexibility exhibited by calreticulin (the chain in purple) and the sparseness of the chain assembly. Fitting the structures to the original experimental map was challenging as indicated by a low TM-Score of 0.5 by VESPER (raw). In contrast, DiffModeler achieved a high TM-Score of 0.95, demonstrating that the diffusion model was effective to capture structural features in the map.

The next map is an example of a complex with many chains (Fig. 3d). It is the state 2 of a complex of the proteolytic core and the ATPase PAN (proteasome-activating nucleotidase) with 34 chains (EMD-213, resolution: 6.35 Å)^49^. DiffModeler was able to fit most of the subunits correct except for long helical domains locating at the top of the complex in the figure, yielding a TM-Score of 0.97. In comparison, with the original map, VESPER (raw) was only able to fill about 20% of the structure with a TM-Score of 0.20.

The last example (Fig. 3e) is a map from the minor state of T. *thermophilus* enzyme in complex with NADH (EMD-11237, resolution: 6.10 Å)^50^, which includes 15 chains. Fitting subunit structures to the original map was difficult as all the chains are α-helical and hard to distinguish as indicated by a low TM-Score of 0.64 by VESPER (raw). On the other hand, with the advantage of the map diffusion, DiffModeler showed accurate backbone structure tracing with backbone recall of 0.98 and a superior structure alignment with a TM-Score of 0.98 and an RMSD of 2.40 Å.

Fig. 4 illustrates the largest protein complex structure built by DiffModeler. This example is a map for proteasome in complex with ADP-AlFx (EMD-6693)^51^ determined at a 6.30 Å resolution. The complex comprises 47 protein chains totaling 13,462 amino acids. The diffusion model in DiffModeler achieved a 0.92 backbone tracing recall, laying a robust foundation for further protein complex structure modeling. Overall, the modeled complex showed high consistency with the native structure, as evidenced by a TM-Score of 0.94 and a sequence identity of 0.89. When individual chains are considered, 45 chains out of 47 chains were successfully modeled with an average sequence matching of 92.6%. The 45 chains include 17 individual chain structures shown in the figure, which appear in the front view of the complex. The high modeling accuracy is clearly due to the application of the diffusion model to the map, as VESPER (raw) alone only achieved 0.25 TM-Score. In contrast, EMBuild yielded a TM-Score of 0.88 and a sequence identity of only 0.47, which indicates many chains were placed on similar but not correct map regions.

**Fig. 4.**
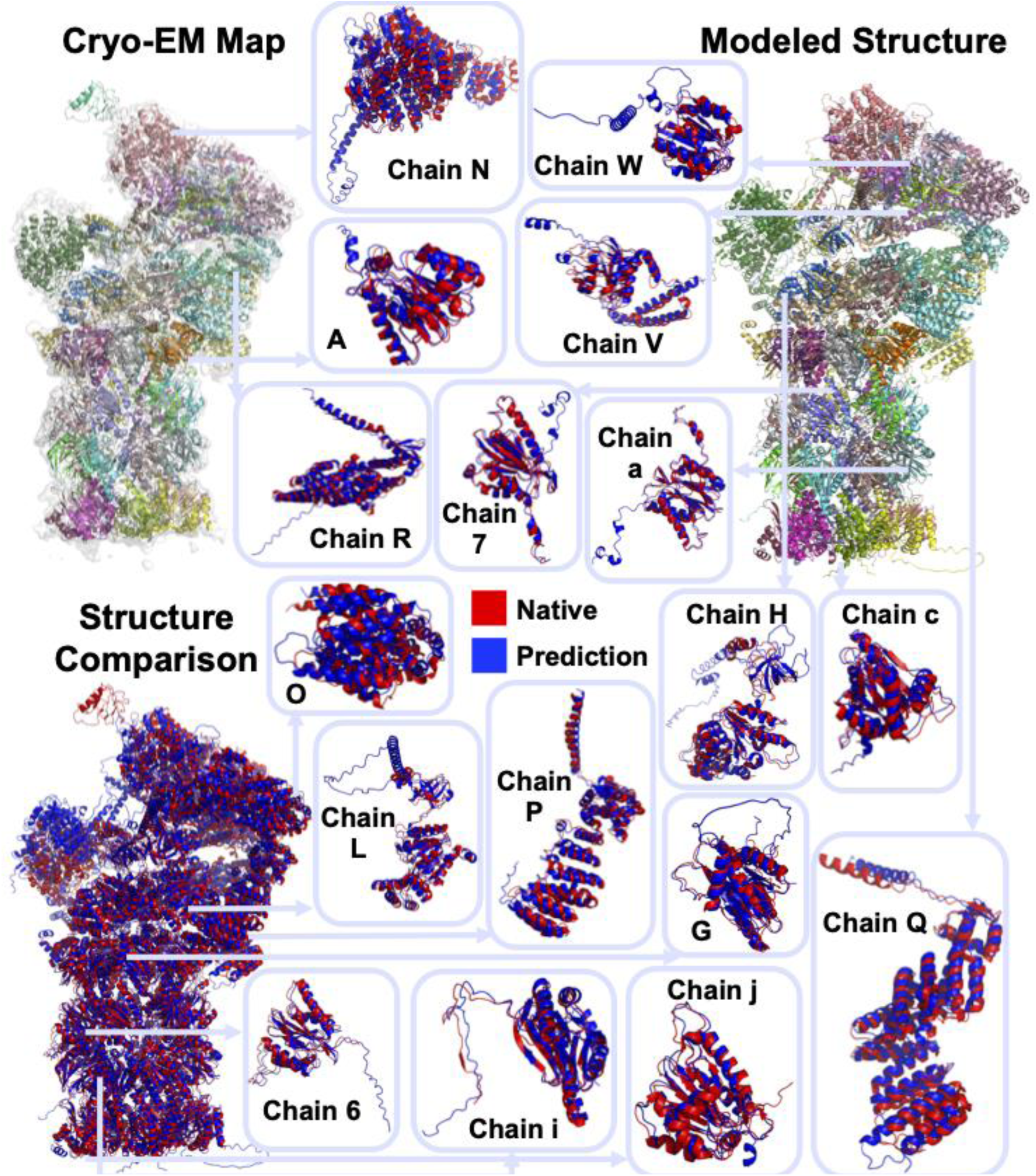
Structure model of proteasome constructed by DiffModeler. This is the largest complex in the test set. Proteasome in complex with ADP-AlFx (EMD-6693, PDB: ID: 5WVI, Resolution: 6.30 Å; 47 chains and 13,462 aa). TM-Score: 0.94, Sequence Identity: 0.89, RMSD: 5.13 Å. The EM map superimposed with the corresponding complex structure and the model by DiffModeler are shown on the top left and top right, respectively, with different colors indicating different chains. On the bottom left, superimposition of the entire complex in PDB and the model is shown together with comparison of 17 individual chain models that appear in the front view of the complex. Blue, the model, red, the native structure in the PDB entry. The modeled structures by the other methods are shown in Extended Data 6.

### Structure Modeling on cryo-EM maps at a lower resolution range

We further conducted an additional benchmark of DiffModeler on cryo-EM maps determined at low resolutions (10 to 18 Å). There were four maps in EMDB, which are in this resolution range and satisfy the map selection criteria we used, i.e. maps with a corresponding PDB entry that have a cross correlation and an overlap higher than 0.65 with the map, and all the chains have an AF2 model that have a TM-Score of 0.5 or higher. The modeling results are shown in Fig. 5 and detailed performance metrics are provided in Supplementary Table 4. For these four maps, the average TM-Score of models by DiffModeler was 0.74, while that of EMBuild, Phenix, and VESPER (raw) was 0.32, 0.36 and 0.27, respectively, which are below the cutoff of 0.5 that indicates meaningful structural similarity.

**Fig. 5.**
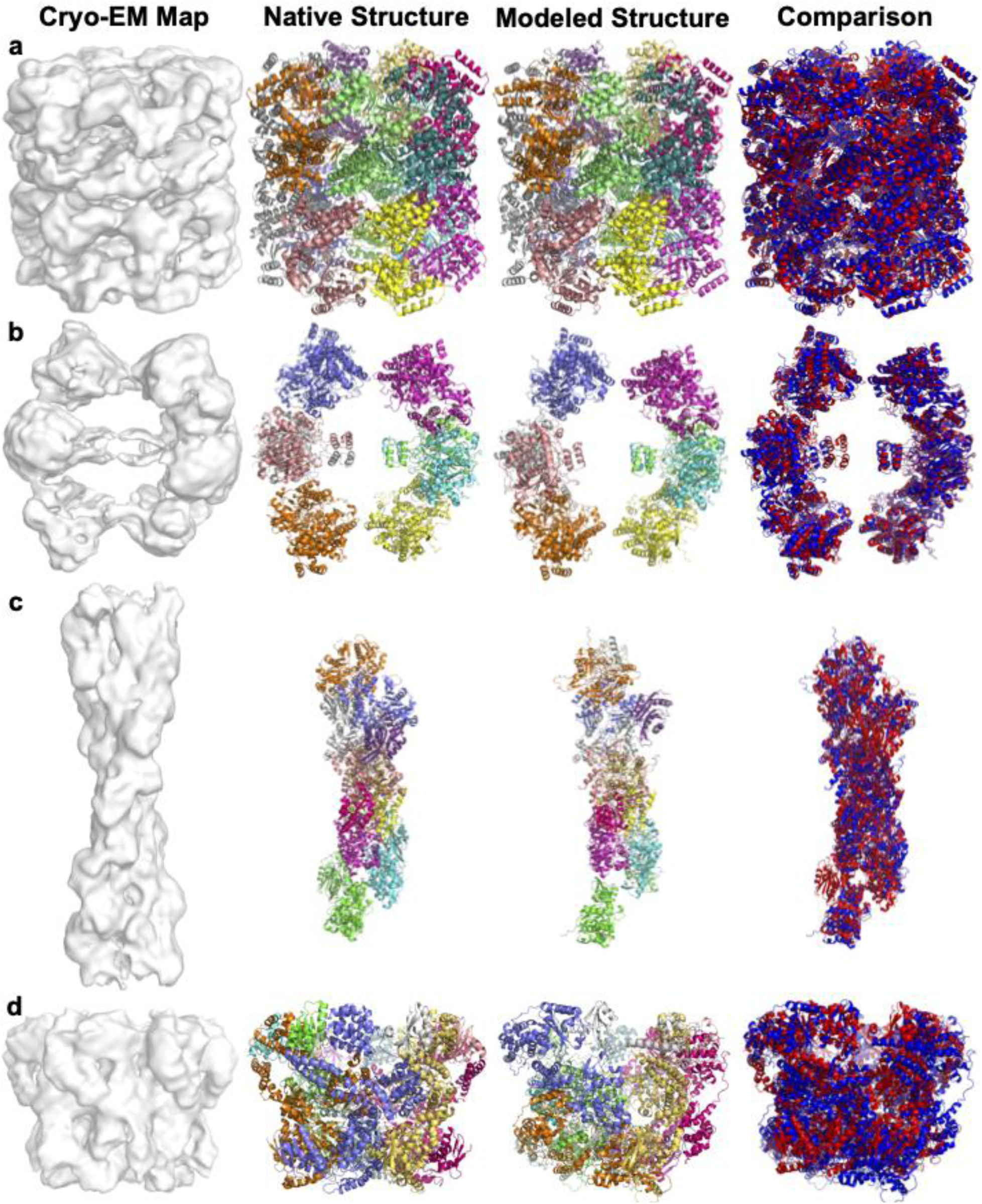
Modeling results by DiffModeler for experimental maps at low resolution. Detailed Evaluation Results are shown in Supplementary Table 4, which contains TM-score of individual chains and modeled protein complex by different methods. For each map four panels are shown from left to right: the input cryo-EM map; the corresponding native structure; the model by DiffModeler; and the superposition of the DiffModeler model (blue) with the native structure (red). **a.** ATP-Bound States of GroEL (EMD-1042, PDB: ID: 1GR5, Resolution: 10.3 Å; 14 chains and 7,238 aa): TM-Score: 0.97, Sequence Identity: 0.95, RMSD: 3.88 Å. **b.** anaerobic fatty acid beta oxidation trifunctional enzyme (anEcTFE) octameric complex (EMD-16134, PDB: ID: 8BNR, Resolution: 10.3 Å; 8 chains and 4,584 aa): TM-Score: 0.87, Sequence Identity: 0.88, RMSD: 5.80 Å. **c.** cofilactin filament inside microtubule lumen (EMD-16877, PDB: ID: 8OH4, Resolution: 16.5 Å; 14 chains and 3,776 aa): TM-Score: 0.60, Sequence Identity: 0.50, RMSD: 8.30 Å. **d**, MecA-ClpC complex with ATP with the Walker B mutations introduced in the D2 ring (EMD-5608, PDB: ID: 3J3S, Resolution: 11.0 Å; 12 chains and 5,352 aa): TM-Score: 0.51, Sequence Identity: 0.45, RMSD: 8.42 Å. Modeling results by other methods are provided in Extended Data 7.

The first example (Fig. 5a) is ATP-Bound States of GroEL (EMD-1042, resolution: 10.3 Å) ^52^, comprising 14 chains with a total of 7,238 residues. Due to the low resolution of the map, the authors manually determined this structure by fitting individual chain structures while considering symmetry information. In contrast, DiffModeler demonstrated the capability to model the complete atomic structure automatically and accurately, achieving a TM-Score of 0.97 and an RMSD of 3.88 Å. The structural superimposition of the model with the corresponding PDB entry visually confirms the accuracy of the model.

The next one is a 10.3 Å map from anaerobic fatty acid beta oxidation trifunctional enzyme (anEcTFE) octameric complex (EMD-16134)^53^. The complex has eight chains, which is a dimer of a tetramer, shown as left and right volumes in the map in the figure. Notably, the original investigators faced challenges in directly resolving the structure at this low resolution. Their strategy involved first elucidating the structure from a tetramer map with a resolution of 3.55 Å. This process utilized fitting and real-space refinement techniques, incorporating the crystal structure information from the PDB entry 6DV2^54^. Subsequently, they docked the solved structure into the low-resolution map and conducted further refinement to achieve the final structure. In marked contrast, DiffModeler automated the assembly of the entire protein complex based on the low-resolution map and achieved a high TM-Score of 0.87. Structures derived from EMBuild, Phenix, and VESPER reported TM-Scores of 0.30, 0.30, and 0.19, respectively, emphasizing the distinct advantage offered by DiffModeler.

The third map was determined even at a lower resolution of 16.5 Å (Fig. 5c). The map contains 14 chains, cofilactin filament inside microtubule lumen (EMD-16877). The authors determined the structure by fitting cofilactin filament model (PDB: 5YU8) to the density manually followed by a local refinement^55^. The model built by DiffModeler had a TM-Score of 0.60, which was substantially higher than values of EMBuild, Phenix, and VESPER (raw), 0.17, 0.25, and 0.18, respectively, which failed to capture even the overall fold. The model by DiffModeler captured the overall shape of the complex. However, only 4 chains out of 14 chains are successfully aligned (sequence identity: 0.99). There were chains, e.g., chains E, H, and K, which were placed in the correct region of the map but with an incorrect alignment.

In the last panel (Fig. 5d), we illustrate a case where DiffModeler’s performance was relatively poor. The presented structure is derived from a 11.0 Å resolution map or a 12-chain complex of MecA-ClpC with ATP and Walker B mutations introduced in the D2 ring (EMD-5608). The authors employed a complex manual procedure for structure determination: Initially, they used an initial model based on another crystal structure of ClpC (PDB: 3PXI) and employed MODELLER^56^ to fill in missing loops using other related structures as templates (PDB: 1JBK and 1R6B). Subsequently, the structure was manually docked into the cryo-EM maps, followed by flexible fitting using NAMD^57^. The model generated by DiffModeler had an overall TM-Score of 0.51, a barely significant score for structure modeling. Among 12 chains, 4 chains C, D, E, and F were modelled successfully with an average TM-score of 0.73 and sequence identity of 0.73. The rest of the chains were placed to incorrect regions of the map. TM-Scores of EMBuild, Phenix, and VESPER (raw) were even worse, 0.26, 0.30, and 0.19, respectively.

To our knowledge, DiffModeler is the first method capable of automatically modeling protein complexes from maps in this low-resolution range. It distinctly demonstrates its advantage over existing methods.

### Structure Modeling for Maps at a Higher Resolution

Although the primary focus of DiffModeler is on maps of low resolution, where it can demonstrate its unique strengths, it also performs effectively with higher resolution maps. To illustrate this versatility, we employed DiffModeler on maps with better than 5 Å resolution. We conducted benchmarking on two distinct datasets: one that was used in the paper of CryoREAD^9^ and the other employed in the work of ModelAngelo^58^. These two datasets cover a broad spectrum of structures, encompassing protein-DNA/RNA complexes and protein-only configurations. The CryoREAD dataset comprised 61 maps (excluding those containing only nucleic acids), while the ModelAngelo dataset included 28 maps. Maps in the datasets varied in terms of the number of protein chains they include, which ranges from 1 to 48 chains. Accordingly, the number of amino acids in maps ranged from 447 to 17,947. On this dataset we used the identical model and pipeline of DiffModeler without any alterations. For maps with protein-DNA/RNA complexes, we first used CryoREAD to construct DNA/RNA structures and then modeled protein structures in the remaining regions in the maps by DiffModeler. AF2 models used in the DiffModeler modeling was selected from the AF2 database by BLAST^59^ sequence search. Supplementary Table 5 shows the sequence identity and the TM-Score of AF2 models relative to the native structure of individual chains in the maps. The TM-Score of AF2 models ranged from 0.134 to 0.998, with an average TM-Score of 0.858 and 0.896 for CryoREAD and ModelAngelo datasets, respectively.

Fig. 6 summarizes the modeling results. Individual modeling results for each map are provided in Supplementary Table 6. In Fig. 6a-d, models of the maps were evaluated with TM-Score and the sequence identity with other methods on the two datasets. To characterize the performance of DiffModeler, three other modeling methods were also employed, Phenix (phenix.dock_in_map), VESPER (raw), and ModelAngelo^58^, a recent deep learning-based modeling method. Among these methods, Diffmodeller clearly outperformed the other three methods on the two datasets. The average TM-Score and the sequence identity by DiffModeler for the CryoREAD/ModelAngelo datasets were 0.879/0.907 (Fig. 6a, 6c) and 0.851/0.864 (Fig. 6b, 6d), respectively, which are comparable to the results obtained for the original dataset of 5.0 to 10.0 Å resolutions as shown in Fig. 2. In contrast, the other three methods showed substantially lower performance: Phenix, VESPER (raw), ModelAngero yielded the average TM-Score, and the sequence identity were 0.572/0.697/0.348 and 0.430/0.579/0.328 on the CryoREAD dataset, 0.573/0.701/0.542 and 0.508/0.605/0.532 on the ModelAngelo dataset. In Fig. 6e, we investigated the impact of complex size on the TM-Score of DiffModeler models. As depicted in Fig. 2, we consistently observed high TM-scores, even for large complexes.

**Fig. 6.**
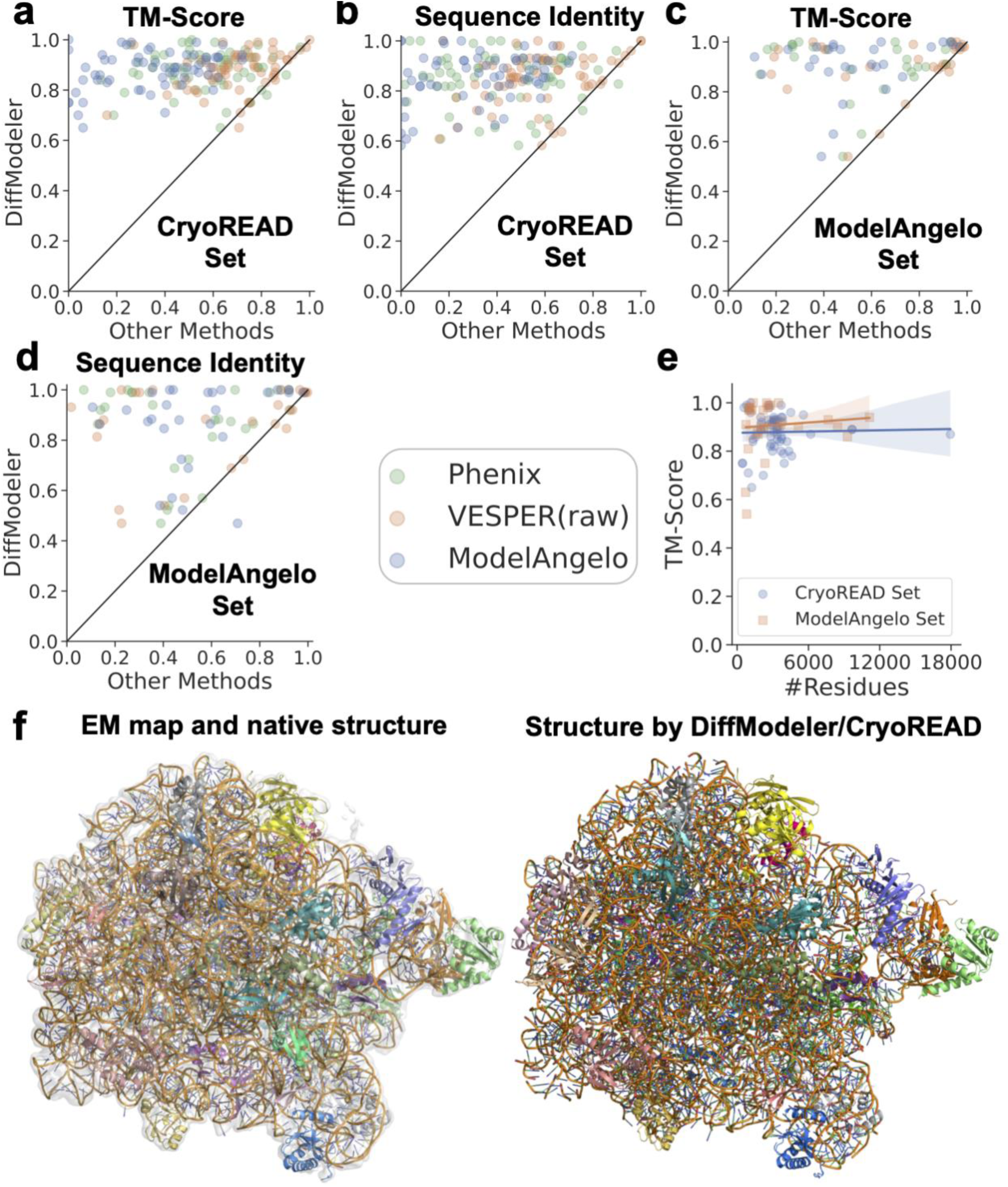
Protein complex structure modeling by DiffModeler for experimental maps at near-atomic resolution (< 5 Å). Detailed Evaluation Results are shown in Supplementary Table 5 and 6. The benchmark was performed on two datasets, the CryoREAD dataset and the ModelAngelo dataset Models by DiffModeler were compared with those by Phenix, VESPER (raw) and ModelAngero. **a.** TM-Score comparison on the CryoREAD dataset. **b.** The sequence identity comparison on the CryoREAD dataset. **c.** TM-Score comparison on the ModelAngelo dataset. **d.** The sequence Identity comparison on the ModelAngelo dataset. **e.** TM-Score relative to the total number of residues in the map. **b.** Sequence identity relative to the total number of residues in the map. **c.** RMSD (Å). **d.** RqcH DR variant bound to 50S-peptidyl-tRNA-RqcP RQC complex (EMD-13017, PDB: ID: 7OPE, Resolution: 3.2 Å; 3,818 residues and 2,996 nucleotides). DiffModeler and CryoREAD: TM-Score: 0.92, Sequence Identity (proteins): 0.92, RMSD: 1.74 Å.

While DiffModeler exhibited strong performance for most cases, there were instances where the performance is low, indicated by TM-Score or sequence identity values lower than 0.6. One contributing factor to these cases was the failure in predicting the AF2 chain structure (e.g., EMD-12935, EMD-27705 (Supplementary Table 6). There are also two cases with a low sequence identity and a high TM-Score (e.g., EMD-13619, EMD-13620), which are cases of hetero-oligomers where subunits were placed in equivalent places of different chains.

In Fig. 6f, we present an example of models by DiffModeler. This model is for a protein-RNA complex of RqcH DR variant bound to 50S-peptidyl-tRNA-RqcP ribosome-associated protein quality-control (RQC) complex (EMD-13017, resolution: 3.2 Å)^60^. This large complex includes 3,818 amino acid residues and 2,996 nucleotides. We modelled the entire complex with DiffModeler and CryoREAD^9^ for protein and RNA regions, respectively, which yielded a TM-Score of 0.92 for the protein region and a backbone recall of 0.94 for the RNA region . While this work primarily focuses on protein structure modeling with DiffModeler, we also extended our modeling efforts to include nucleic acid structures within these maps using CryoREAD^9^. Backbone and sequence recall (the fraction of sugar and phosphate atoms that were placed within 5 Å and the fraction of nucleotides that were correctly identified in the model), were measured at 0.855 and 0.523 on the CryoREAD dataset and 0.829 and 0.413 on the ModelAngelo dataset.

## Discussion

DiffModeler is a novel structure modeling method, which uniquely targets low resolution cryo-EM maps of 5-15 Å. Within this target resolution range, accurate modeling of protein structure complexes presents significant challenges. Not only *de novo* structure modeling^4,5,37,58^, fitting known structures to the map is also difficult as presented in the results with VESPER (raw). The presence of noisy density in cryo-EM maps makes it exceedingly difficult to detect precise atom and amino acid positions as well as main-chain conformations in the map. DiffModeler overcomes these obstacles by sculpting out main-chain conformations from low resolution maps using a diffusion model and by representing the salient points with LDPs, which enables to achieve substantially higher accuracy in structure fitting. The benchmark of DiffModeler on higher resolution, better than 5 Å, further indicated its generalizability and accuracy to handle maps with higher resolution. DiffModeler may appear similar to existing deep learning-based methods^61,62^, which modifies or sharpens maps, but it is distinct because it is not simply for sharpening map density; rather, it is a multi-step pipeline that outputs complex structure model as the end product.

Although DiffModeler has demonstrated overall accuracy and effectiveness, it is crucial to address the limitations of the current version. First, in some regions with low local resolution, the backbone tracing of diffusion model may be inaccurate, leading to incorrect structure fitting. To address this issue, further enhancements can be made to prioritize the fitting of regions with higher local resolution, mitigating the risk of such errors. Secondly, as of now, DiffModeler exclusively supports protein structure complex modeling. To expand its applicability, future developments will aim to extend its capabilities to support protein/DNA/RNA complex structure modeling, enhancing its versatility in addressing a wider range of biological systems. Furthermore, for high-resolution cryo-EM maps (better than 4 Å), it will be essential to develop local structure refinement approaches that leverage the density information to refine predicted structures, further enhancing accuracy and reliability. Addressing these limitations remains as future developments.

We firmly believe that DiffModeler will prove to be an indispensable and user-friendly tool for protein complex structure modeling, bridging a crucial gap in the availability of tools suitable for maps at low resolutions. The approach will also be applicable for cryo-electron tomography within the same resolution range, better than 15 Å, which is now increasingly available^63,64^.

## Acknowledgments

The authors thank Jacob C. Verburgt, Anika Jain, Charles Christoffer for their help in literature search, discussion, and proofreading. The author would also thank Jessica A. Nash, Sam Ellis and Jing Chen’s suggestion for optimizing the released software.

## Author contributions

DK conceived the study. XW designed and implemented DiffModeler and computed results. HZ and GT optimized the VESPER algorithm and participated in implementing the full pipeline. All the authors analyzed the results. XW drafted the manuscript and DK edited it. All the authors read and approved the manuscript.

## Materials & Correspondence

Daisuke Kihara

## Code availability

The source code of DiffModeler is made available at https://github.com/kiharalab/DiffModeler. It can run on our webserver https://em.kiharalab.org/algorithm/DiffModeler freely without installing it in a local machine. We also provide sequence version of DiffModeler on our server https://em.kiharalab.org/algorithm/DiffModeler(seq), which can automatically use the sequence information to find the most similar single-chain structure from RCSB and AlphaFold database and then model the full protein complex structure. The source code of ComplexModeler (including DiffModeler and CryoREAD) for protein-DNA/RNA complex structure modeling is made available at https://github.com/kiharalab/ComplexModeler. It is also available our webserver https://em.kiharalab.org/algorithm/ComplexModeler.

## Data availability

The entries of the maps and corresponding structure models utilized in this study are provided in Supplementary Tables 1, 4 and 6. The experimental EM maps utilized can be downloaded from the EMDB (https://www.emdataresource.org/). The corresponding experimental determined structures utilized can be downloaded from the RCSB (https://www.rcsb.org/). The structures modeled by DiffModeler and the corresponding native structures from RCSB are available at https://doi.org/10.5281/zenodo.10714883.

## Methods

### Constructing the benchmark dataset

Following the protocols employed in our previous works^19,20,37,65^, we complied a dataset of experimental cryo-EM maps for training, validation, and testing DiffModeler. Initially, we sourced cryo-EM maps from EMDB (as of January 26^th^, 2023) with resolutions between 5 Å to 10 Å and had the corresponding deposited structures in PDB with more than 20 residues. We only kept maps that contain only proteins. This initial screening yielded 840 maps.

Subsequently, we assessed the quality of structure-to-map fit by measuring cross-correlation and overlap between the EM maps and simulated maps generated from their respective structures in PDB ^17^. Maps were discarded if their corresponding structures displayed a cross correlation and overlap below 0.65. The remaining maps were manually inspected. These steps reduced the number of maps to 337.

To remove redundancy in the data, we applied single linkage clustering with the sequence identity of proteins within each map. Two maps were grouped into the same group if any protein chains from both maps exhibited a global sequence identity of 25% or higher. This clustering procedure resulted in 103 clusters. Out of the 103 clusters, we randomly allocated 68 clusters (230 maps) for the training set, 18 clusters (36 maps) for validation, 17 clusters (71 maps) for testing (Supplementary Table 1). It is important to note that the training, validation, and testing sets are fully independent from each other. Finally, we further filtered maps in the testing set that contained inaccurate predicted models with a TM-score lower than 0.5 in the Alphafold Database^18^. This filtering process resulted in the final testing dataset comprising 19 maps, reduced from 71 maps.

### Pre-processing of map data

If a map had a grid size that is different from 1.0 Å, we interpolated the grid size to 1.0 Å using trilinear interpolation. The density values within a map were normalized to [0.0, 1.0] with a minimum-maximum normalization. Any negative values in a map were set to 0, and 0 was used as the minimum value for normalization. We set the maximum value for normalization as the 98th percentile density value, and any density values above that were capped at 1.0.

From each map, boxes of a size of 64^3^ Å^3^ were collected by scanning the box across a map along three axes with a stride of 32 Å. Each grid point within the box was assigned a label indicating whether it belonged to the backbone. If a grid point was within 2.0 Å of any backbone atoms, it was assigned as backbone. Otherwise, the point was considered as background. A box was excluded from training if less than 0.1% of the grid points were assigned as backbone.

### Training the conditional diffusion model of DiffModeler

Given the density information from cryo-EM maps, the objective of the diffusion model of DiffModeler is to generate the backbone labels in the map. We employed a conditional diffusion model, particularly, the denoising diffusion implicit model (DDIM)^66^, for its superior generation quality and efficiency. Inspired by the Pix2Seq^28^ framework, we designed an encoder-decoder network architecture (Extended Data 1). The encoder scans the input density map with a box of 64^3^ size and embeds (outputs) hidden features of the map. The decoder utilizes three components as input of the conditional diffusion framework: the condition (the starting cryo-EM density map and hidden features), the noised backbone 𝑥_𝑡_ at timestep *t*, and the time *t* of the current step. From these inputs, the decoder outputs the predicted traced backbone 𝑦_𝑡_. The noised backbone 𝑥_𝑡_ is a mixture of the ground-truth traced backbone density 𝑥_0_and the Gaussian noise 𝜀 determined by the timestep *t*, which will be explained later. The encoder and the decoder are optimized simultaneously by comparing the predicted traced backbone 𝑦_𝑡_ and ground truth traced backbone 𝑥_0_. The encoder and decoder neural network architecture is shown in Extended Data 2a and 2b, respectively. The detailed network architecture of each component of the encoder and the decoder is shown in Extended Data 3.

As mentioned in the previous dataset section, we allocated 230 maps for training and 36 maps for validation of the conditional diffusion model. For each batch of training, we randomly sampled 8 boxes from the 230 maps. In total, around 16,000 and 3,500 boxes were used in an epoch for training and validation, respectively. The framework was trained through 30 epochs, and the final model is selected based on the validation performances.

The main objective of the model is to perform conditional denoising of a noisy density of the traced backbone to achieve the ground-truth traced protein backbone density *x0* in the map. For training the model, a series of noisy traced backbone density maps were generated by randomly sampling the density values from the ground-truth traced backbone density and the Gaussian noise:

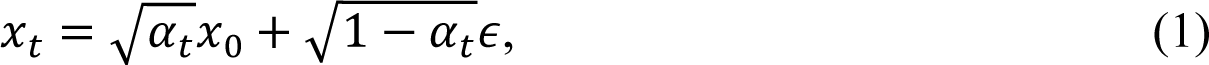

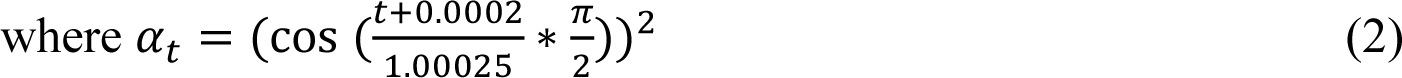

where 𝑥_𝑡_ is the noised traced backbone at timestep 𝑡, 𝛼_𝑡_ is a cosine scheduling function shown in Eq. (2), 𝑥_0_ is the ground-truth traced backbone, and 𝜖 is a noise variable randomly sampled from the standard Gaussian noise, 𝒩(𝟎, 𝑰). The ground-truth density of the traced protein backbone was prepared by assigning the backbone label to each grid point based on the corresponding backbone native structure (N, Cα, C atoms). For any grid point in the map, if a grid point was within 2.0 Å of any backbone atoms, it was assigned as backbone. Otherwise, the point was considered as background.

During the training process, 𝑡 was uniformly sampled from [0,1] for each map in the training set at each iteration to enforce that the framework successfully captures the diffusion process. The noised backbone 𝑥_𝑡_ for time *t* was obtained according to Eq. (1), from which the decoder computes the predicted backbone map 𝑦_𝑡_. The loss of 𝑦_𝑡_ was computed in comparison with the ground truth backbone 𝑥_0_. The used Dice loss^36^ was define as

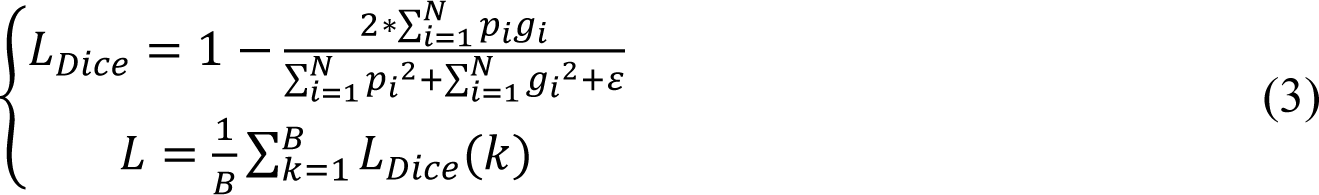

𝐿_𝐷𝑖𝑐𝑒_ represents the Dice loss of a predicted box 𝑃 of prediction 𝑦_𝑡_ at timestep 𝑡 and a corresponding ground truth box 𝐺 of ground truth 𝑥_0_; 𝑁 is the total number of grid points inside the box; 𝑝_𝑖_ ∈ 𝑃 is the predicted probability of the *i*-th grid point in the predicted box; 𝑔_𝑖_ ∈ 𝐺 is the binary ground truth of the *i*-th grid point, where 1 denotes the existence of backbone structure in the grid point and 0 indicates background; 𝜀 is a smoothing factor with value of 1e-6; 𝐿 is the overall loss of a batch of 𝐵 examples; 𝐿_𝐷𝑖𝑐𝑒_(𝑘) represents the dice loss of *k*-th example’s detection. Here different samples in the same batch may have different timestep 𝑡 since it is uniformly and independently sampled for each example.

We tested hyperparameter combinations of a learning rate of [1e-3, 1e-4, 1e-5] with a weight decay of [0, 1e-6, 1e-5, 1e-4] using the Adam optimizer ^67^. Among the combinations, the learning rate 1e-4 without weight decay showed the best grid-wise Intersection-over-Union (IoU) of 0.562 on the validation set. Training and validation of the conditional diffusion model took around 5 days. The computations are performed on two paralleled NVIDIA RTX A6000 48 GB GPU connected via NVLink.

### Inference of the conditional diffusion model in DiffModeler

With the trained conditional diffusion model, we compute the traced backbone conditioned on the input cryo-EM density. The inference of conditional diffusion model is presented in Extended Data 4. Given a box of cryo-EM density, the encoder of the conditional diffusion model first embeds the hidden features of the input density box. Subsequently, the decoder starts with the random Gaussian noise as the initial distribution 𝑥_𝑇_ and iteratively refines the estimated density from 𝑡 = 𝑇 to 𝑡 = 0 to make it closer to the ground-truth traced backbone 𝑥_0_, conditioned on the hidden features from the encoder and the initial density input.

Benefited from the training, which used uniformly sampled timesteps, we have the flexibility to choose the overall inference steps *T*. We chose T = 100 as we did not observe significant performance improvement with *T* larger than 100. The current timestep *t* is calculated by

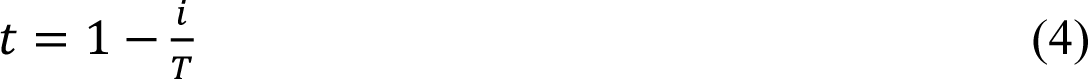

where 𝑡 is the timestep at inference iteration 𝑖, and 𝑇 is the overall inference steps.

The first iteration of the inference starts at timestep T. The decoder takes the random Gaussian noise 𝑥_𝑇_, timestep 𝑇 embedding, and the condition (i.e., the hidden feature embedding and the original cryo-EM map) as input and then it outputs 𝑦_𝑇_.

In the following iterations with timestep 𝑡 = 𝑇 − 1, 𝑇 − 2,…,0, the condition inputs are the same and the timestep 𝑡 embedding obtained with Eq. (4). However, the noisy backbone input 𝑥_𝑡_for decoder is different from training. During training, *xt* is computed following Eq. (1) which uses 𝑥_0_as the ground-truth traced backbone. As 𝑥_0_ is not available in the inference stage, the input of the decoder, 𝑥_𝑡_, uses the decoder’s output 𝑦_𝑡+1_ at timestep 𝑡 + 1:

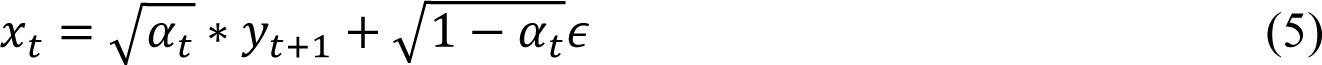

where 𝑥_𝑡_ is the estimated noised backbone at timestep 𝑡, 𝑦_𝑡+1_ is the decoder output at 𝑡 + 1 and 𝜖 is the random Gaussian noise. In this equation, 𝜖 is also estimated by comparing the decoder’ noisy backbone input 𝑥_𝑡+1_ and its corresponding backbone estimation output 𝑦_𝑡+1_ from the decoder as follows:

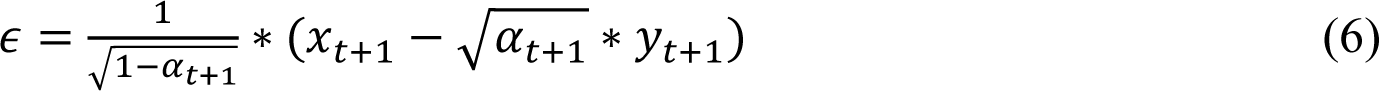

By combining Eq.(5) and Eq.(6), we can obtain the decoder input 𝑥_𝑡_ with decoder output 𝑦_𝑡+1_ at timestep 𝑡 = 𝑇 − 1, 𝑇 − 2,…,0. The inference process of the decoder is repeated for 𝑇 = 100 times and 𝑥_0_ at timestep 𝑡 = 0 is our final estimated backbone.

### Single-chain structure fitting using VESPER

We used VESPER^16^ for fitting AF2 models of individual proteins to the modified map by the diffusion model. AF2 models of the protein chains were taken from the Alphafold database^18^. Supplementary Table 3 provides TM-score of the chains. The average TM-score was 0.922. The fitting process involved three main steps: Initially, AF2 models were transformed into simulated maps at a 1 Å resolution using TEMPy^68^. In the subsequent step, we simplified both the modified EM map and the simulated maps of the AF2 models into maps by condensing them into maps with local representative density points. This was achieved through the mean-shifting algorithm^41^ a method we devised in our early work, MAINMAST^4^. Finally, VESPER was used to globally align AF2 models into various poses within the representative map, generating different fit scores. The top 100 poses were retained as pose candidates for each subunit.

The mean shift algorithm is employed to compute maps featuring local representative density points by clustering density points within an EM map. First, grid points with a density exceeding 0 are identified. Then, the algorithm iteratively updates the coordinates of a grid point 𝑥 by considering the weights associated with neighboring grid points: 𝑥^𝑡+1^ = 𝑓(𝑥^𝑡^), where

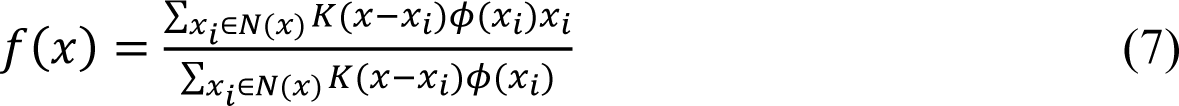

𝑁(𝑥) is the neighborhood of 𝑥, which are a set of neighboring grid points that satisfy ||𝑥_𝑖_ − 𝑥||_2_ ≤ 2 ∗ 𝜎; 𝐾(𝑝) is a Gaussian kernel function with bandwidth 𝜎, as shown in Eq.(8); 𝜙(𝑥) is the density value of the grid point 𝑥.

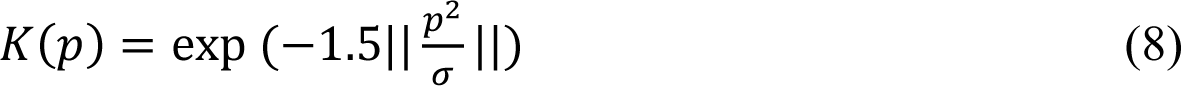

where the 𝜎 is the bandwidth set as 2. The mean-shift process is continued until convergence, i.e., ||𝑥^𝑡+1^ − 𝑥^𝑡^||_2_ ≤ 𝛿 with 𝛿 set to 0.001.

Following the completion of the mean-shifting process, we merged shifted points that were in close proximity. Points closer than a predefined threshold distance of 2.0 Å, were clustered together, and the grid point with the highest density within the cluster was designated as the representative node. This clustering and selection process was iterated until the convergence of the selected representative nodes. The resulting set of points, known as representative points, forms the basis for the representative map (Fig. 3).

By completing this stage, we acquired two distinct representative maps using the mean-shift algorithm: the subunit representative map (𝑅𝑀_𝑠𝑢𝑏𝑢𝑛𝑖𝑡_) derived from the simulated map of the AF2 single-chain structure, and the backbone representative map (𝑅𝑀_𝑏𝑎𝑐𝑘𝑏𝑜𝑛𝑒_) obtained from the diffusion-traced backbone map.

The final step involves utilizing VESPER to globally align AF2 single-chain subunits into various poses within the backbone map. Specifically, VESPER aligns different 𝑅𝑀_𝑠𝑢𝑏𝑢𝑛𝑖𝑡_ to 𝑅𝑀_𝑏𝑎𝑐𝑘𝑏𝑜𝑛𝑒_ obtained in the preceding step. For each subunit representative map 𝑅𝑀_𝑠𝑢𝑏𝑢𝑛𝑖𝑡_ , VESPER systematically explores all potential poses to align 𝑅𝑀_𝑠𝑢𝑏𝑢𝑛𝑖𝑡_ with 𝑅𝑀_𝑏𝑎𝑐𝑘𝑏𝑜𝑛𝑒_ . In VESPER’s global search, we used a rotation scan interval of 10 ° and a translation scan interval of 2 Å. The fitness score of 𝑅𝑀_𝑠𝑢𝑏𝑢𝑛𝑖𝑡_ *i* at pose *j* is defined as:

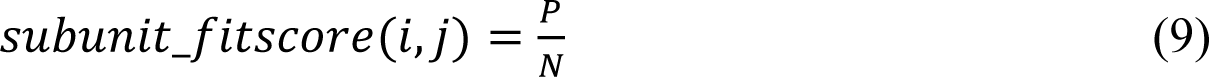

where *P* is the number of Cα positions of subunit 𝑖 at pose 𝑗 that have representative points in 𝑅𝑀_𝑏𝑎𝑐𝑘𝑏𝑜𝑛𝑒_ within 3 Å, and 𝑁 is the total number of Cα positions of subunit 𝑖. Top 100 poses were kept for each subunit. This pool comprises of 𝑀 ∗ 100 pose candidates for a protein structure complex with 𝑀 chains.

### Assembling subunits to generate the entire protein complex structure

Subunits, fitted to the map with different pose candidates, are then assembled into a complete protein complex structure model. We developed a greedy algorithm that iteratively assembles superimposed subunits within the map. The entire pipeline is depicted in Extended Data 5. As outlined in the preceding section, we generated 100 poses for each subunit in the map using VESPER. Therefore, the subunit-pose pool for a given protein structure complex comprises *M*100* pose candidates, all of which were scored using the *subunit_fitscore* (Eq. 9).

The initial step in the modeling process involves selecting the subunit-pose with the highest *subunit_fitscore* among all available poses. Subsequently, a local region within 20 Å from the fitted subunit-pose is masked out in the backbone map 𝑅𝑀_𝑏𝑎𝑐𝑘𝑏𝑜𝑛𝑒_ and the subunit pose is further optimized in terms of the *subunit_fitscore* with an interval of 5° for rotation scan and an interval of 1 Å for translation scan in that local region. Then, from the subunit-pose pool, subunit-poses are removed if the poses belong to the subunit that was just selected or if they have significant overlap with the selected subunit-pose. A subunit-pose is considered to have overlap if more than 10% of Cα positions of the subunit-pose are closer than 3 Å to any Cα positions to an already selected subunit-pose(s).

Following this, the subsequent best subunit-pose is selected iteratively until the subunit-pose pool is exhausted. In most cases, where each subunit assumes a correct pose, all M subunits are successfully fitted into the map. However, there are rare instances where not all subunits are selected due to significant overlap among all 100 poses of a subunit with other already-selected subunit-poses. In such scenarios, where some subunits remain unfitted due to substantial overlap, a new pose set is generated for these remaining subunits. This is achieved by fitting them to the remaining density regions within 𝑅𝑀_𝑏𝑎𝑐𝑘𝑏𝑜𝑛𝑒_ using VESPER. The same iterative process is then applied until all the subunits are successfully fitted.

---

**Extended Data 1.**
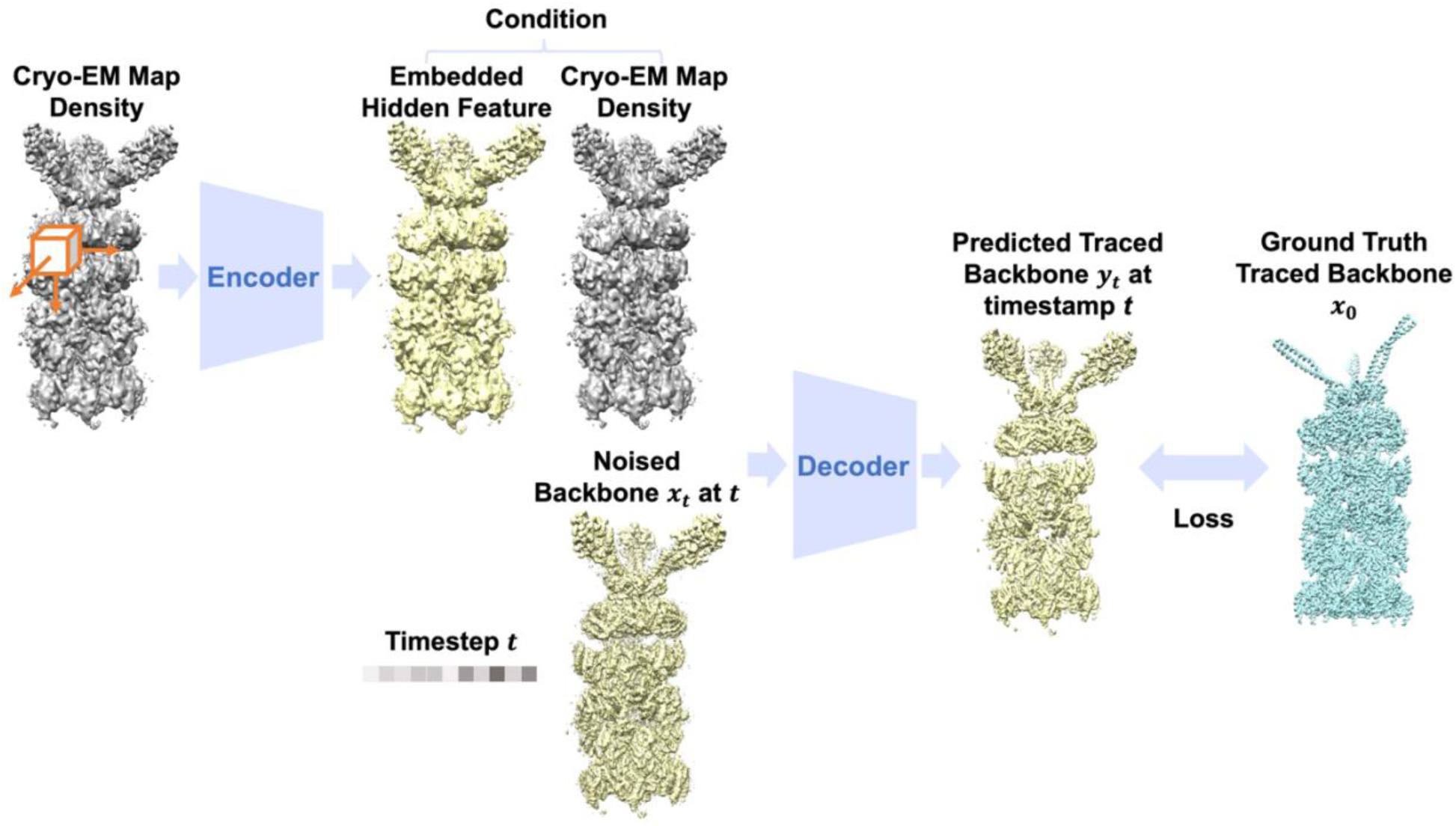
The overall framework of the conditional diffusion model in DiffModeler. The entire framework consists of one encoder and one decoder. The encoder takes the cryo-EM density as input and outputs the hidden features by scanning the map density with a box. The decoder utilizes three main components as the input of the conditional diffusion framework: the condition (the starting cryo-EM density map and hidden features), the noised backbone 𝑥_𝑡_ at timestep *t*, and the timestep *t*. Then the decoder outputs the predicted traced backbone 𝑦_𝑡_.The encoder and the decoder are optimized simultaneously by comparing the predicted traced backbone 𝑦_𝑡_ and ground truth traced backbone 𝑥_0_, with details illustrated in Methods.

**Extended Data 2.**
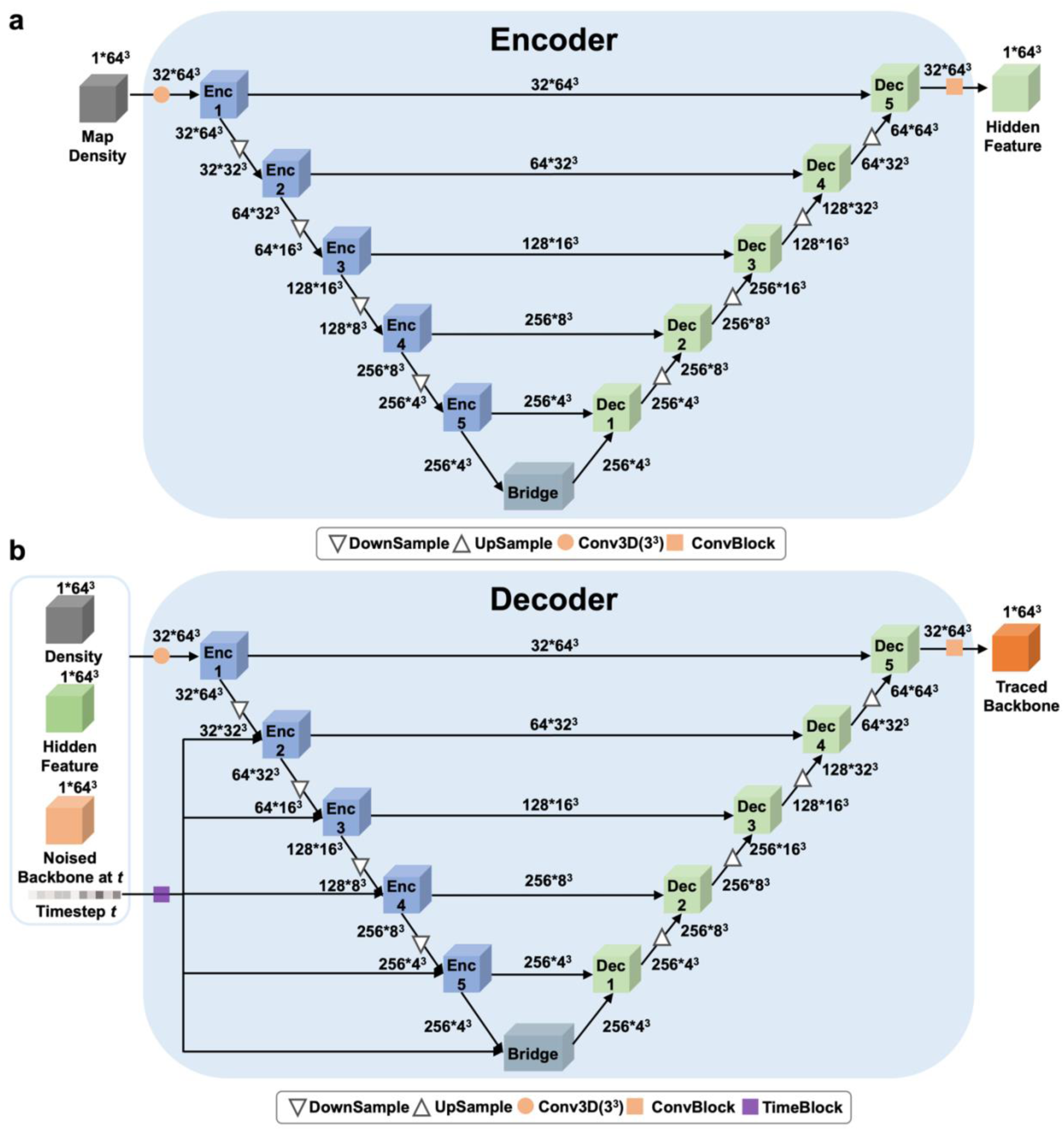
The network architecture of the conditional diffusion model in DiffModeler. **a.** The encoder network architecture. It is a 3D U-shape-based convolutional neural network (UNet) with skip connections. The channel size of different layers is also illustrated in the figure. The input is first processed by Conv3D layer with 32 filters in size of 3^3^, and then iteratively processed and down-sampled by encoding block Enc1-Enc5 (Extended Data 3a), The downsample block (Extended Data 3c), the dense information is further processed by bridge block (Extended Data 3b), subsequently process the encoding, which is upsampled by Dec1-Dec5 (Extended Data 3a). The upsample blocks (Extended Data 3d) with skip-connections connecgting with the encoding blocks, and the final ConvBlock(Extended Data 3e) aggregate the information and yield the final output. **b.** The decoder network architecture. It shares a similar UNet architecture as the encoder. Additionally, it includes a TimeBlock (Extended Data 3f) that encodes the timestep input and passes it to every levels of encoding block in the decoder network. Individual blocks are illustrated in Extended Data 3.

**Extended Data 3.**
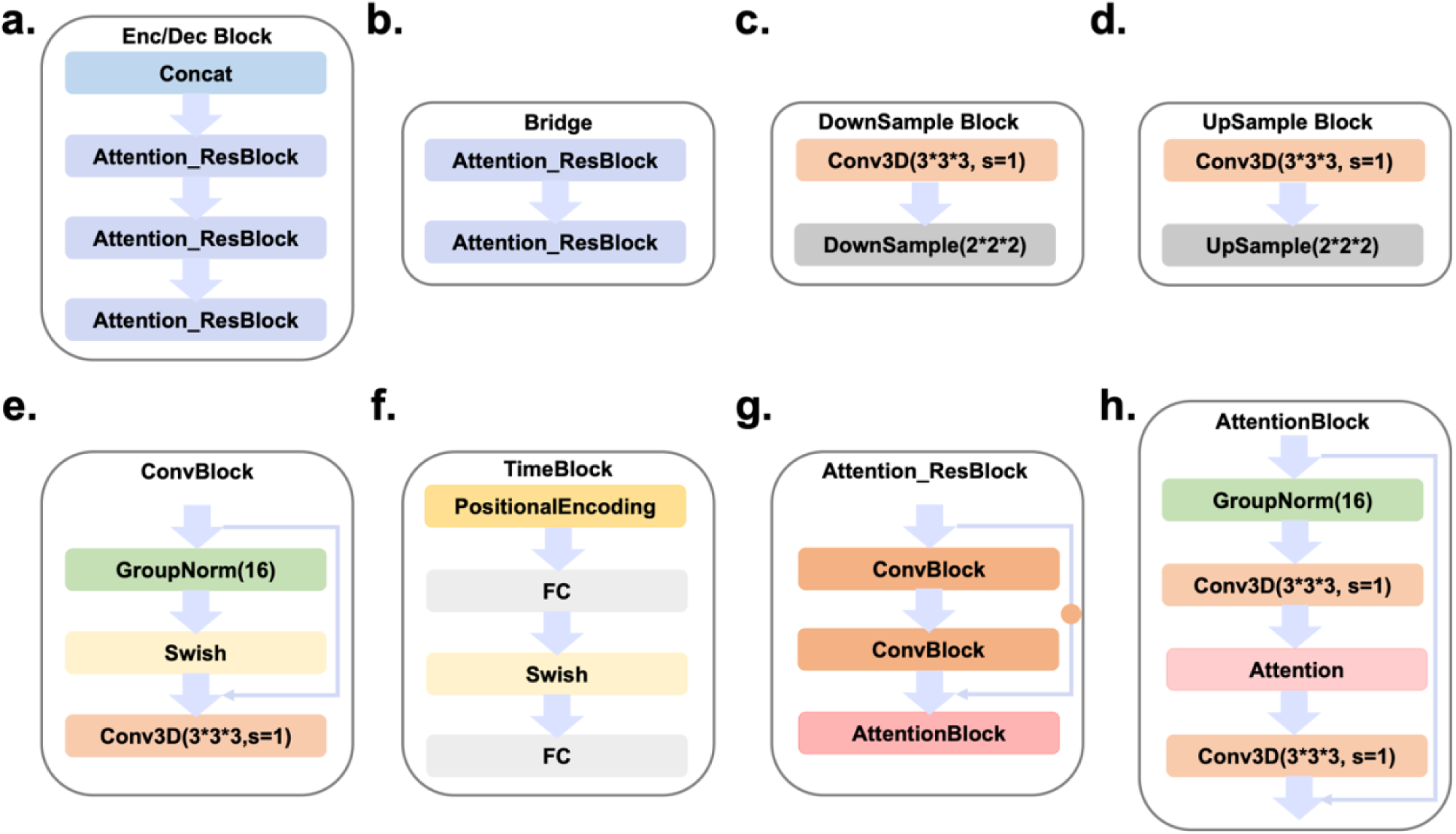
Individual network block architecture of the conditional diffusion model in DiffModeler. **a.** The encoder/decoder block (Enc1-Enc5, Dec1-Dec5. in panel a and b of Extended Data 2). Concat is an operation that concatenates inputs. **b.** The bridge block (located at the bottom of Extended Data 2a, 2b); **c.** The DownSample Block. Conv3D is a 3-dimentional (3D) convolutional layer with a filter size of 3*3*3, stride 1, and padding 1. **d.** The UpSample Block. **e.** The ConvBlock (located one step before the output box in Extended Data 2a, 2b). GroupNorm^69^ is a normalization layer that calculates group statistics across channels to normalize the input data by dividing multiple channels into different groups. Swish^70^ is a smooth, non-monotonic function that consistently matches or outperforms ReLU and serves as an activation layer. **f.** Time Block, specifically designed for timestep embedding. PositionalEncoding^71^ is an explicit layer with pairs of sine and cosine functions to add positional information to the input. FC is a fully connected layer in which each neuron applies a linear transformation to the input vector through a weight matrix. **g.** Attention_ResBlock (Attention_ResBlock in panel a, b); **h.** The Attention Block (AttentionBlock in panel g). Attention^71^ is a layer that enables to dynamically highlight the relevant features of the input data through the attention mechanism.

**Extended Data 4.**
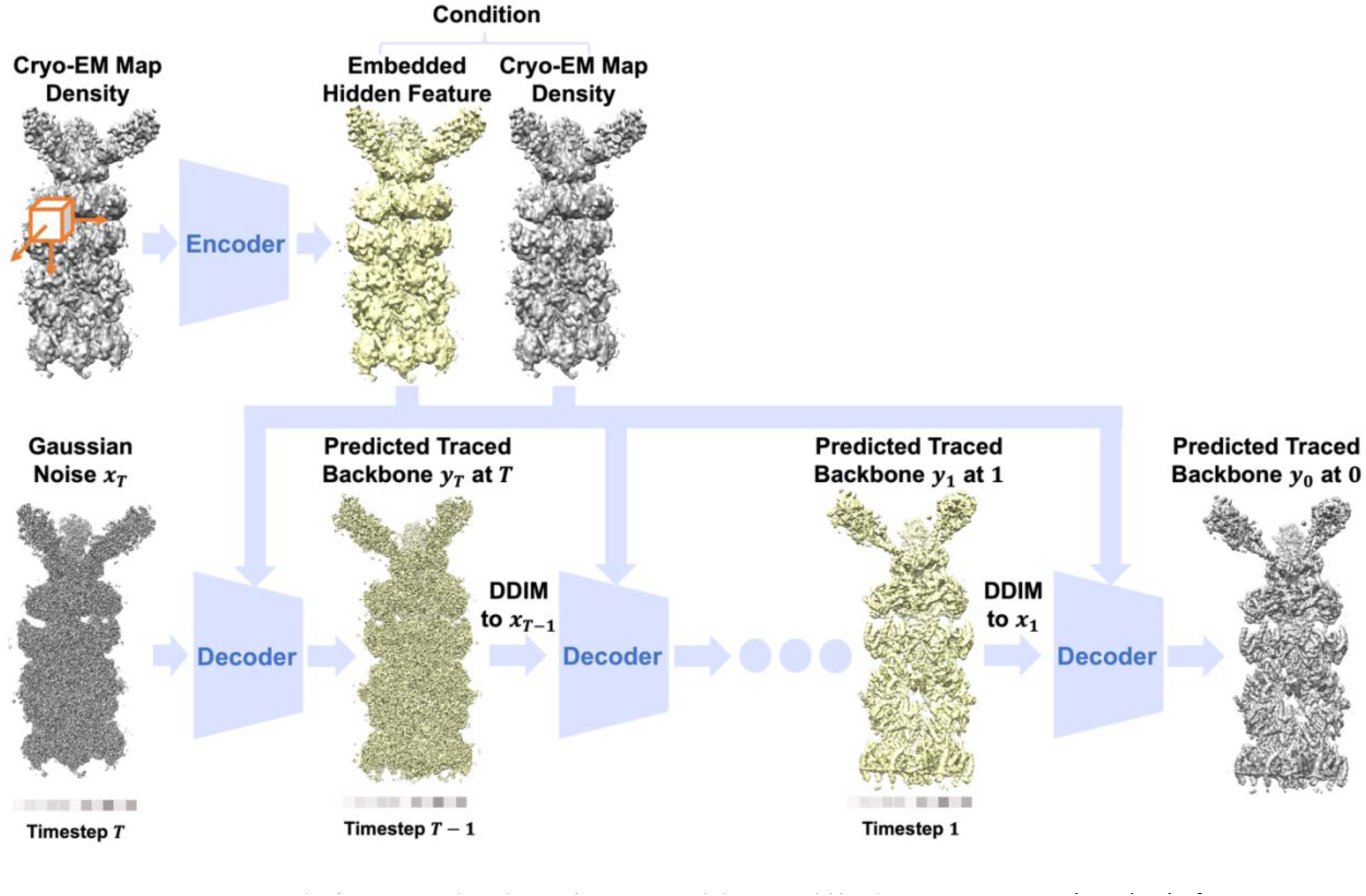
The inference pipeline of the conditional diffusion model. During the inference stage, the encoder first takes the cryo-EM density as input and outputs the hidden features. Then, the decoder iteratively refines the density from time step T utilizing three core information as input to estimate the traced backbone: condition (cryo-EM density and hidden features), the noised traced backbone at timestep *t*, the embedding of timestep t. The noised traced backbone starts with a random Gaussian noise 𝑥_𝑇_ and 𝑥_𝑡_ at timestep 𝑡 is iteratively updated by the decoder’s output 𝑦_𝑡+1_ through DDIM step (illustrated in Methods).

**Extended Data 5.**
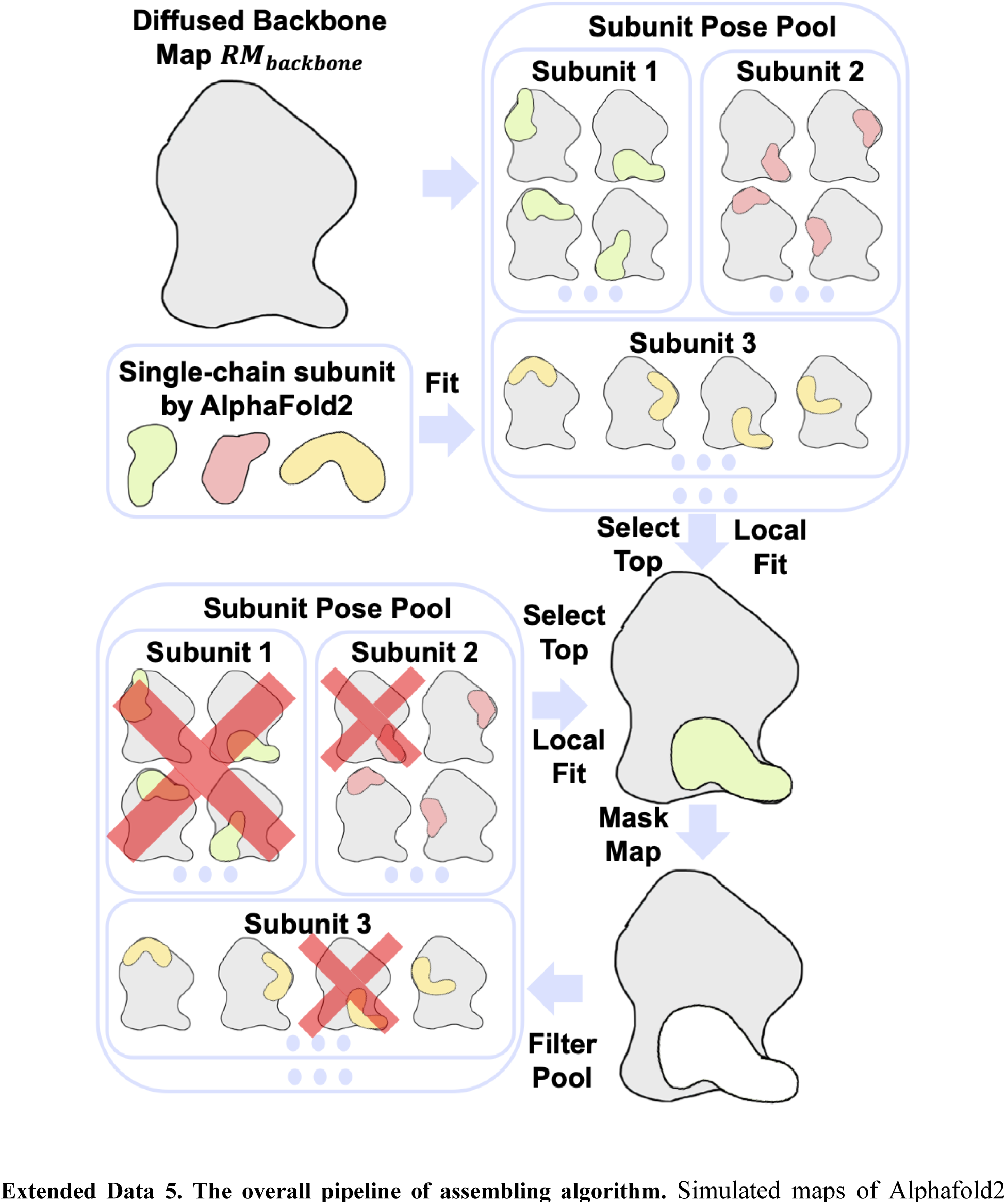
The overall pipeline of assembling algorithm. Simulated maps of Alphafold2 models for each subunit are aligned with the RM_backbone using VESPER, and the top 100 poses for each subunit are cataloged in the structure pose pool. Initially, the subunit-pose exhibiting the highest *subunit_fitscore* is chosen from the structure pool. The local density region within the map occupied by this subunit is then masked out (the white local region in the figure). The pose of the selected subunit undergoes further refinement by VESPER, employing smaller angle and shifting intervals. Subsequent subunit-poses in the pool are eliminated under two conditions as shown by red crosses in the figure: if they belong to the same subunit as the one just selected or if they overlap with the selected subunit-pose. This iterative process continues until all the subunits are chosen to construct the full complex, at which point the pool becomes empty.

**Extended Data 6.**
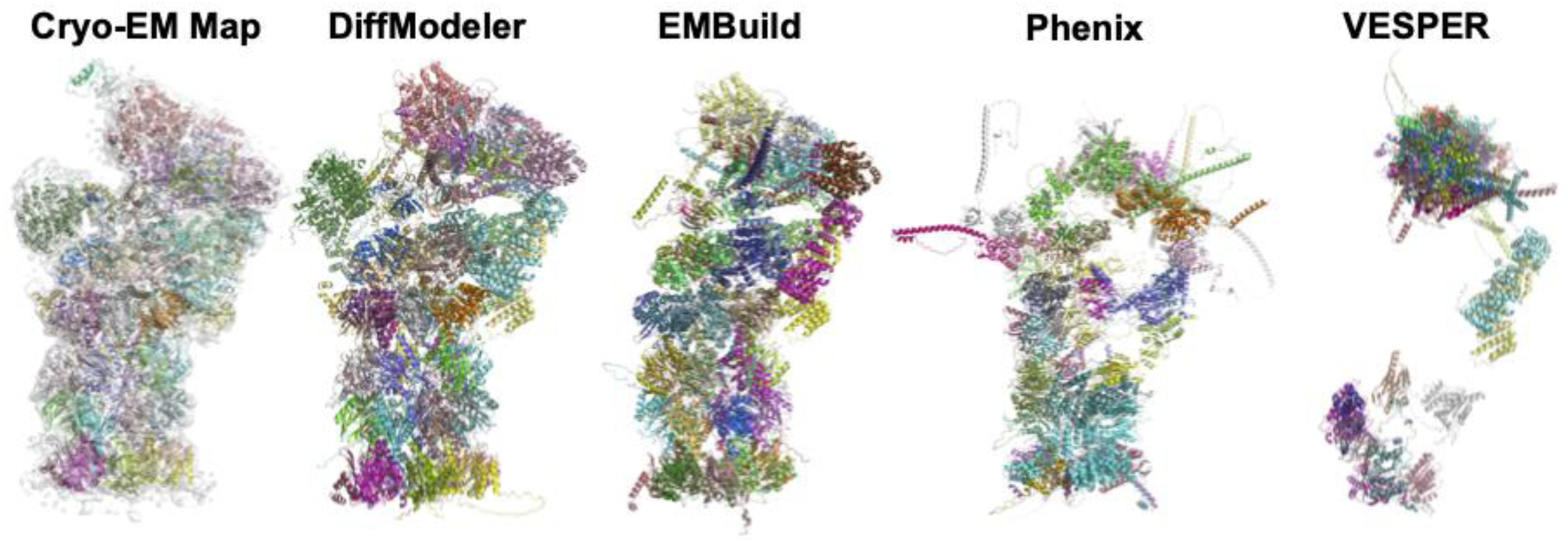
Atomic structure modeling by different methods for experimental map EMD-6872. The proteasome in complex with ADP-AlFx (EMD-6693, PDB: ID: 5WVI, Resolution: 6.30 Å; protein lengths: 47 chains and 13,462 amino acids (aa)). The 5 columns from left to right are 1) EM map and its corresponding structure; 2) the atomic structure by DiffModeler: TM-Score: 0.94, Align Ratio: 0.96, Sequence Identity: 0.89, RMSD: 5.13 Å; 3) the atomic structure by EMBuild: TM-Score: 0.88, Align Ratio: 0.91, Sequence Identity: 0.47, RMSD: 4.8 Å. 4) the atomic structure by Phenix: TM-Score: 0.38, Align Ratio: 0.51, Sequence Identity: 0.04, RMSD: 17.4 Å; 5) the atomic structure by VESPER: TM-Score: 0.25, Align Ratio: 0.29, Sequence Identity: 0.16, RMSD: 12.5 Å.

**Extended Data 7.**
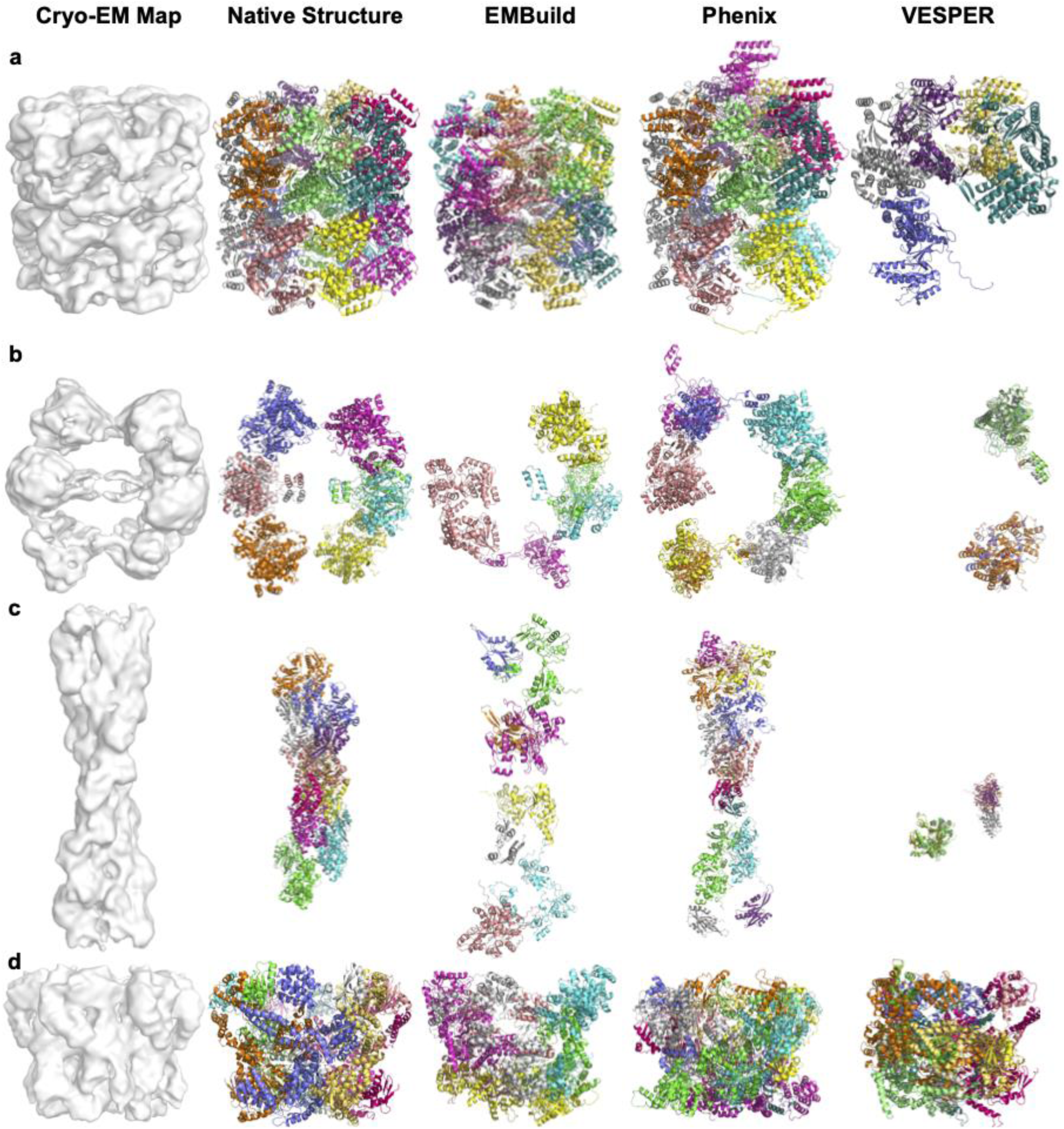
Examples of structure models built with three other methods for experimental maps at low resolution (10-15 Å). Detailed Evaluation Results are shown in Supplementary Table 4. In each row of the modeling example, five columns shown from left to right are 1) input cryo-EM map; 2) the corresponding native structure; 3) the structure model by EMBuild; 4) the model by Phenix; 5) the model by VESPER (raw). The DiffModeler model is shown in Fig. 6. **a**. ATP-Bound States of GroEL (EMD-1042, PDB: ID: 1GR5, Resolution: 10.3 Å; protein lengths: 14 chains and 7,238 amino acids (aa)): DiffModeler (EMBuild, Phenix, VESPER): TM-Score: 0.97 (0.96 0.78, 0.37), Sequence Identity: 0.95 (0.92, 0.71, 0.34), RMSD: 3.88 Å (4.90 Å, 6.50 Å, 7.76 Å). **b**. acid beta oxidation trifunctional enzyme (anEcTFE) octameric complex (EMD-16134, PDB: ID: 8BNR, Resolution: 10.3 Å; protein lengths: 8 chains and 4,584 aa): DiffModeler (EMBuild, Phenix, VESPER): TM-Score: 0.87 (0.30, 0.30, 0.19), Sequence Identity: 0.88 (0.23, 0.17, 0.16), RMSD: 5.80 Å (9.65 Å, 11.28 Å, 8.03 Å). **c**. cofilactin filament inside microtubule lumen (EMD-16877, PDB: ID: 8OH4, Resolution: 16.5 Å; protein lengths: 14 chains and 3,776 aa): DiffModeler (EMBuild, Phenix, VESPER): TM-Score: 0.60 (0.17, 0.25, 0.18), Sequence Identity: 0.50 (0.02, 0.05, 0.08), RMSD: 8.30 Å (12.1 Å, 13.19 Å, 12.67 Å). **d**, MecA-ClpC complex with ATP with the Walker B mutations introduced in the D2 ring (EMD-5608, PDB: ID: 3J3S, Resolution: 11.0 Å; protein lengths: 12 chains and 5,352 aa): DiffModeler (EMBuild, Phenix, VESPER): TM-Score: 0.51(0.26, 0.26, 0.48), Sequence Identity: 0.45(0.16, 0.15, 0.41), RMSD: 8.42 Å(12.28 Å, 11.91 Å, 8.72 Å).

