## Supplemental Data 1 for "DiffModeler: Large Macromolecular Structure Modeling in Low-Resolution Cryo-EM Maps Using Diffusion Model"

**Supplementary Table 1.** (in a separate Excel file)

**Experimental maps in the DiffModeler benchmark dataset.** The file includes four sections. The first three sections are "training", "validation", and "testing", which provide information of clustered EM maps in the training, the validation, and the testing set, respectively. The 1st column, "Cluster" indicates the cluster ID. The 2nd column, "PDB-ID (EMD-ID)", shows the structures that belong to the cluster with their PDB ID and EMD ID. The fourth section "benchmark" shows the map information of maps used to benchmark the DiffModeler. "EMD-ID" is the EMD ID of cryo-EM map, "PDB-ID" is the PDB ID of corresponding protein structure, "Resolution" column shows the reported resolution of the deposited map, "Contour" is the author recommended contour level for the map.

**Supplementary Table 2.** (in a separate Excel file)

**Backbone tracing performance of diffusion model in DiffModeler for individual maps.** "origin map" section includes the grid-level tracing performance from original map, "Diffusion processed map" section provides grid-level tracing performance from the diffusion model. The EMD-ID column represents each entry's EMD ID. "PDB-ID" is the PDB ID of corresponding protein structure. "Resolution" column shows the reported resolution of the deposited map. "Recall" indicates the fraction of correctly identified grid points of backbone, where the denominator is the total number of the backbone grid points from the reference structures (i.e. the PDB entry). On the other hand, "Precision" provides the fraction of correctly predicted grid points, where the denominator is the total number of predicted grid points of backbone.

**Supplementary Table 3.** (in a separate Excel file)

**Atomic structure evaluation for cryo-EM maps in the testing set of different methods.** The "DiffModeler", "EMBuild", "VESPER" and "Phenix" sections provide the performance of DiffModeler, EMBuild, VESPER and Phenix, respectively. The "AlphaFold-Single-Chain" section provides the performance of AlphaFold predicted single chain structure. These tables include all the information of Fig. 2.

The EMD-ID, and PDB-ID columns represent each entry's EMD ID and PDB-ID, respectively. The Resolution column shows the reported resolution of the deposited map. #Chain and #Res indicate the number of protein chains and residues in the deposited structure, respectively. TM-Score reports the TM-Score metric between the predicted structure and native structure. "Align Ratio" quantifies the fraction of residues in the native structure that are successfully aligned by MM-align with the predicted structure. "SeqID" is the fraction of residues in the native structure that are successfully aligned to residues in the predicted structure that share the same residue type, where the denominator is the number of residues in the native structure. "SeqID(Align)" only measures the sequence identity in the aligned residues, where the denominator is the number of aligned residues in the native structure. RMSD is the root-mean-squared-deviation of aligned residues between predicted structure and native structure. "Chain_ID" indicates the specific chain ids in the corresponding structure and "Uniprot_ID" is the chain's corresponding Universal Protein Resource Identifier to each protein entry in the UniProt database.

**Supplementary Table 4.** (in a separate Excel file)

**Atomic structure evaluation for cryo-EM maps in the low resolution dataset of different methods.** The "DiffModeler", "EMBuild", "VESPER" and "Phenix" table provides the performance of DiffModeler, EMBuild, VESPER and Phenix, respectively. The "AlphaFold-Single-Chain" tables provides the performance of AlphaFold predicted single chain structure. These tables correspond to Fig. 5.

The EMD-ID, and PDB-ID columns represent each entry's EMD ID and PDB-ID, respectively. The Resolution column shows the reported resolution of the deposited map. #Chain and #Res indicate the number of protein chains and residues in the deposited structure, respectively. TM-Score reports the TM-Score metric between the predicted structure and native structure. "Align Ratio" quantifies the fraction of residues in the native structure that are successfully aligned by MM-align with the predicted structure. "SeqID" is the fraction of residues in the native structure that are successfully aligned to residues in the predicted structure that share the same residue type, where the denominator is the number of residues in the native structure. "SeqID(Align)" only measures the sequence identity in the aligned residues, where the denominator is the number of aligned residues in the native structure. RMSD is the root-mean-squared-deviation of aligned residues between predicted structure and native structure. "Chain_ID" indicates the specific chain ids in the corresponding structure and "Uniprot_ID" is the chain's corresponding Universal Protein Resource Identifier to each protein entry in the UniProt database.

**Supplementary Table 5.** (in a separate Excel file)

**AlphaFold2 single-chain structure evaluation for the near-atomic resolution datasets.**

The EMD-ID, and PDB-ID columns represent each entry's EMD ID and PDB-ID, respectively. The Resolution column shows the reported resolution of the deposited map. "Chain_ID" indicates the specific chain ids in the corresponding structure and "AFDB_ID" is the chain's corresponding Identifier to single-chain protein entry in the AlphaFold database. TM-Score reports the TM-Score metric between the AF2 predicted structure and native structure. "SeqID" is the fraction of residues in the native structure that are successfully aligned to residues in the predicted structure that share the same residue type, where the denominator is the number of residues in the native structure. RMSD is the root-mean-squared-deviation of aligned residues between predicted structure and native structure.

**Supplementary Table 6.** (in a separate Excel file)

**Atomic structure evaluation for cryo-EM maps in the near-atomic resolution datasets of different methods.** The "DiffModeler(CryoREAD data)" indicates the performance DiffModeler method on CryoREAD dataset, the "DiffModeler(ModelAngelo data)" indicates the performance DiffModeler method on ModelAngelo dataset. We also include tables of other methods on those two datasets: ModelAngelo, VESPER(raw), Phenix. These tables correspond to Fig. 6.

The EMD-ID, and PDB-ID columns represent each entry's EMD ID and PDB-ID, respectively. The Resolution column shows the reported resolution of the deposited map. #Res and #Nuc indicate the number of residues and nucleotides in the deposited structure, respectively. TM-Score reports the TM-Score metric between the predicted protein structure and native protein structure. "Align Ratio" quantifies the fraction of residues in the native structure that are successfully aligned by MM-align with the predicted protein structure. "SeqID" is the fraction of residues in the native structure that are successfully aligned to residues in the predicted protein structure that share the same residue type, where the denominator is the number of residues in the native structure. "SeqID(Align)" only measures the sequence identity in the aligned residues, where the denominator is the number of aligned residues in the native protein structure. RMSD is the root-mean-squared-deviation of aligned residues between predicted protein structure and native protein structure. "Backbone_Recall" quantifies the fraction of backbone atoms of DNA/RNA that were closer than 5 Å to the atoms in the matched nucleotide, where the denominator is the total number of the atoms the reference structures. Similarly, "Sequence_Recall" measures the fraction of correctly identified nucleotides with correct base type assignments, where the denominator is the total number of the nucleotides from the reference DNA/RNA structures.
